# Overt visual attention and value computation in complex risky choice

**DOI:** 10.1101/2020.12.08.416313

**Authors:** Xinhao Fan, Jacob Elsey, Aurelien Wyngaard, Youping Yang, Aaron Sampson, Erik Emeric, Moshe Glickman, Marius Usher, Dino Levy, Veit Stuphorn, Ernst Niebur

## Abstract

Traditional models of decision making under uncertainty explain human behavior in simple situations with a minimal set of alternatives and attributes. Some of them, such as prospect theory, have been proven successful and robust in such simple situations. Yet, less is known about the preference formation during decision making in more complex cases. Furthermore, it is generally accepted that attention plays a role in the decision process but most theories make simplifying assumptions about where attention is deployed. In this study, we replace these assumptions by measuring where humans deploy overt attention, *i.e.* where they fixate. To assess the influence of task complexity, participants perform two tasks. The simpler of the two requires participants to choose between two alternatives with two attributes each (four items to consider). The more complex one requires a choice between four alternatives with four attributes each (16 items to consider). We then compare a large set of model classes, of different levels of complexity, by considering the dynamic interactions between uncertainty, attention and pairwise comparisons between attribute values. The task of all models is to predict what choices humans make, using the sequence of observed eye movements for each participant as input to the model. We find that two models outperform all others. The first is the two-layer leaky accumulator which predicts human choices on the simpler task better than any other model. We call the second model, which is introduced in this study, TNPRO. It is modified from a previous model from management science and designed to deal with highly complex decision problems. Our results show that this model performs well in the simpler of our two tasks (second best, after the accumulator model) and best for the complex task. Our results suggest that, when faced with complex choice problems, people prefer to accumulate preference based on attention-guided pairwise comparisons.

## 1 Introduction

The essence of decision-making is selecting between two or more alternatives, each typically defined by several attributes. An informed choice is necessarily preceded by collecting information about attributes of one or more of the alternatives. The properties of a (partial or complete) subset of attributes of a (partial or complete) subset of alternatives can either be compared directly and conclusions then drawn about the relative merits of the associated alternatives, or they can be integrated to allow comparison between alternatives, after which the subjectively most attractive alternative will be selected. In the latter case, formal measures of attractiveness in this context go back to the period of enlightenment. One possibility is the expected value [13], another the expected utility [2, 48].

### 1.1 Direct value assignment to alternatives

These methods, as well as the more recent Prospect Theory [43, 18] which builds on them, are agnostic about how the gathering and processing of information is performed. Known limitations of cognitive capabilities of humans and other animals strongly suggest that a parallel, simultaneous processing of all necessary information is impossible in all but the simplest situations. The evaluation process must therefore be at least partially of a sequential character. If it is possible to determine the desirability of all alternatives by taking into account all of their attributes, a simple and efficient algorithm to make use of all available information is as follows. Assuming we can integrate all attributes of a given alternative to obtain its overall value, we perform this action for one of the alternatives (which may be chosen randomly or by some other criterion) and store its value together with (a pointer to) its identity in memory. The same is done for another alternative and the two values are compared. The higher (better) of these values replaces the originally stored value and the identity of the alternative with the higher value is stored alongside. Then another, not yet evaluated alternative is selected and the process repeated. After all alternatives have been evaluated this way, the best alternative (the one with the highest value) is the one stored in memory. This algorithm scales linearly with the product of the number of alternatives and attributes. Furthermore, it has low storage requirements: one storage “slot” for the highest value found so far, another for the identity of the alternative corresponding to it, plus one slot for the identity of the alternative currently under evaluation and one for its value. Additional memory capacity may be required for keeping track of which alternatives have been evaluated. This can be as little as one memory slot if the alternatives are evaluated in a fixed sequence, *e.g.* by going down the list of alternatives from top to bottom.

While this algorithm is efficient and seemingly frugal in resources, there are many empirical results indicating that in many cases, this is not the method chosen by humans and other animals. One piece of evidence is the existence of preference reversal behavior [49, 39, 40, 12] that is incompatible with such an algorithm. Deciders thus must use different strategies, possibly because the process of integrating attribute properties to associate a value to each alternative, which is essential for the described algorithm, is in many cases not possible or not easy.

### 1.2 Comparisons between attributes

It has been argued [46, 44] that it is easier to establish categorical differences (*e.g.* “this attribute is better than that one”) than to determine the relative value of one alternative *vs.* another. Indeed, there are decision procedures that do not require such value assignment. Instead, they rely on comparisons between attribute values rather than integration of all attribute values for each alternative. An influential early method is elimination by aspects [42]. Here, it is assumed that deciders rank the attributes (called “aspects” in the original publication) in the order of their importance. Starting with the most important attribute, alternatives considered inferior with respect to this attribute are eliminated from the competition. For those remaining, the second-most important attribute is considered and again the least preferred alternatives are eliminated. This procedure continues until one alternative remains which is then chosen. Only relative comparisons within one attribute are required in this algorithm, together with an elimination threshold. Even simpler is a lexicographic rule in which the highest ranked candidate in the most important attribute is selected, except in a tie in which the next-most important attribute is chosen. Tversky [41] showed that even in this simple situation intransitivity of choices (and thus preference reversals) can occur if insensitivity to just-noticeable-differences is allowed.

Another example of deciding by within-attribute comparisons is Priority Heuristic [5]. In this model the decider goes through a series of heuristic rules designed to minimize losses and stops when the result is “good enough,” again without ever integrating attributes to determine alternative values. A categorical selection rule which does not rely on outright elimination of alternatives by single attributes is Decision By Sampling [38, 35]. Pairs of attribute values from different alternatives are randomly selected from all available ones. The alternative with the more favorable attribute is upvoted by one unit. Instead of eliminating alternatives outright based on a single comparison, an alternative is selected if the tally of its votes (from all of the sampled attributes) exceeds a pre-determined threshold, at which point the competition is terminated. This model also includes a long-term memory component that we do not discuss here.

### 1.3 Complex decision problems

Much of the work discussed so far has been concerned with relatively simple decisions in laboratory settings. The field of Operations Research (or Management Science) deals with problems of multi-attribute decision making (called Multi Criteria Decision Making, or MCDM in this field) whose complexity often exceeds by far that of assays used in psychological research. As in the simpler situations discussed above, also for these more complex scenarios assigning quantities like utility functions to each alternative may be much more difficult than making pairwise comparisons of attributes. To systematically combine the results of these comparisons, “outranking” methods were developed in Operations Research in which relative rankings of alternatives are established systematically based on the pairwise comparisons between attributes [31].

A prominent example of outranking methods is the Preference Ranking Organisation METHod for Enrichment Evaluation (PROMETHEE) [6, 7]. In the first step of this method, pairwise differences between values of the same attribute in all alternatives are tabulated (a 2-dimensional array, see eq. 23 below). These attributes can be of very different character, therefore the PROMETHEE method transforms these differences into a common (scalar) representation space. The mapping between attribute differences and the common representation can be done by means of different classes of functions. These “preference functions” can be continuous or discrete, piece-wise linear or nonlinear but they all have in common that as function of the value difference they (i) vanish at zero and below zero, (ii) reach unity for the maximum difference, and (iii) monotonically increase between these extremes. Examples of such functions are given in the cited references; we use a piece-wise linear function in section 2.9. In our extension of the model which we introduce in section 2.10 to take into account the attentional state of the deciders, we introduce our own advantage function. In particular, our functions deviates from condition (i) discussed above, *i.e.* our function can take on negative values, see Figure 4.

The weighted sum over all the preferences of all attributes of one alternative over another is the overall preference of the first alternative over the second. Entering this preference as one entry in another two-dimensional array whose indices are the two alternatives, we see that the sum over all entries in row *i* in this array is the overall advantage that alternative *i* has over all other alternatives (the “positive outranking flow” of alternative *i*). Conversely, the sum over all entries in column *i* is the overall advantage that all other alternatives have over alternative *i* (“negative outranking flow” of alternative *i*). The ranking of alternative *i* is the difference between these two flows, and in the version of the model that we use in sections 2.9 and 2.10 (PROMETHEE II; [6]), the highest-ranked alternative is predicted to be the one that is selected by the decider.

### 1.4 Attentional dynamics of decision making

It is highly likely that complex decisions are made through a sequence of steps rather than instantaneously. Our goal is to understand the process of decision making and we surmise that following the steps taken should improve understanding of the process over simply looking at the final outcome. A classical method is a verbal protocol in which deciders give a “running commentary” of their perception of their decision process, either concurrently with the decision process or retroactively. Limitations are that both types of protocols may interfere with the ongoing task, are likely incomplete, are difficult to analyze, and yield only approximate information about the timing of different stages of the decision process. One alternative to observe the process of information acquisition is using the Mouselab environment [16, 27] in which deciders use a computer mouse to indicate which attributes they choose to be displayed on a computer screen (more recent versions are web-based, see mouselabweb.org). A more direct approach is to monitor eye movements during the decision process [32, 33, 10, 37, 19, 9]. Advantages of this approach are that data are easy to collect, quantify and analyze, that data collection interferes only to a minimal extent with natural behavior, and that there is a rich literature concerned with the role of eye movements in other fields, *e.g.* perception [23, 24]. For these reasons, this is the approach we adopt for the behavioral work of the present study, section 2.1.

We suggest that observing eye movements does not only tell us about sequentiality of decision making processes but that the insight they provide is more profound. Eye movement are closely linked to the attentional state of the observer and several models of decision making have an attentional component in which alternatives in the space of potential choices are selected, *e.g.* [8, 30, 17, 3]. However, the attentional state of the decider is typically assumed to be not known and various assumptions are made, *e.g.* random jumps between possible selections. Eye movements (*i.e.* overt attention) provide an estimate of the attentional state of humans (and other animals). In the field of perceptual attention, the predictive power of a computational model is typically quantified by comparing its predictions for the location of the focus of attention with fixations made by human (and simian [1]) observers that free-view static or dynamic scenes [25, 4]. This is possible because, even though it has been known for more than a century that covert attention can be dissociated from overt attention [47], the latter is often a good approximation of the former^1^. This, then, makes it possible to go beyond making assumptions of the attentional state during the decision making process and replace them by actually measuring the state.

We describe how we measure the eye movements of healthy volunteers while they perform decision making tasks of varying complexities in section 2.1. The main part of this report is devoted to the development of models that predict choice behavior, sections 2.2-2.10. In other words, the main purpose of this study is to explore the predictive power that overt attention (eye movements) has on the eventual decision of a human decider. We develop several models of the influence of overt attention and we compare their predictive power, both between the different models and with “baseline models” that do not take into account eye movements.

## 2 Results

### 2.1 Human behavior during multi-attribute decision making with known attentional state

We study the behavior of human participants in a task in which they choose one alternative out of several available. Each alternative is defined by multiple attributes. We investigate two cases that differ substantially in complexity. In the simpler case (“2×2”), two alternatives are presented with two attributes each while in the more complex situation (“4×4”), there are four alternatives with four attributes each. Obviously, for an exhaustive evaluation of all aspects a minimum of four attributes needs to be assessed in the first task while the more complex one requires 16. Alternatives are presented on a computer screen and, importantly, the values of all attributes are by default covered by opaque circles whose colors indicate the *type* of an attribute but not its *value*. The value is “hidden” under the opaque disks and only is replaced by a symbol indicating the value of the corresponding attribute upon active fixation of the observer. This is accomplished by the use of an eye tracker that continuously monitors the eye position of the head-fixed observer and permits attribute value unmasking within a few tens of milliseconds, making the switch barely noticeable. Disk colors are consistent for all experiments: yellow for amount to win, blue for probability to win, red for amount to lose (only for 4×4), and green for the delay until feedback becomes available (*ditto*). For details of the experimental paradigm see section 4.

Figure 1a shows the stimulus configurations used in the experiment. Participants could collect all information they desired by looking selectively at those attributes they were interested in at a given point in time, as many times as they desired, until they indicated the choice of their preferred alternative by pressing the associated key on a keyboard. Figure 1b shows eye traces during example trials. Figure 1c is an overview of all phases of one trial, starting when the participant directed their gaze at a fixation spot, then at a series of locations representing the 2 attributes of the available 2 alternatives in this particular task, and terminating with the selection of one of the alternatives by pressing a key on a keyboard.

**Figure 1:**
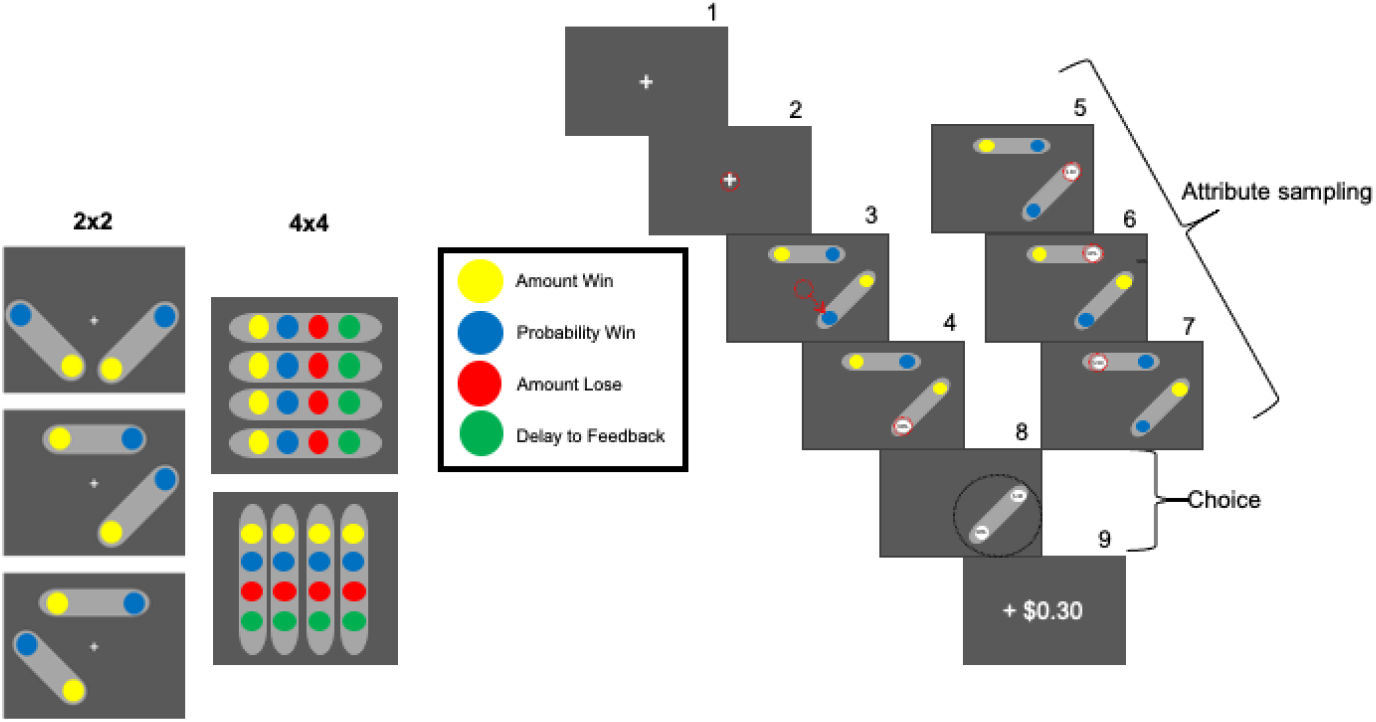
Task design. (**a**) Choice menu layout and attribute types. Left: Examples of two-alternative, two-attribute (2×2) configurations, Right: Examples of four-alternative, four-attribute (4×4) configurations. (**b**) Task flow for one example 2×2 trial.

In a forthcoming publication (Elsey *et al.*, in preparation) different attribute sampling strategies used by the participants will be analyzed in detail; for simple examples see the two strategies illustrated in Figure 1b. In the current study, we instead investigate the effectiveness of predicting human behaviors in this task for a wide range of computational models, ranging from entirely parameter-free to some with about a dozen free parameters. Specifically, we want to predict the choices humans make in non-trivial tasks, defined as those trials in which no alternative is better in one attribute than in all others and not worse than in all other attributes. Thus, a compromise between attributes is required that depends on the individual preferences of the participants, there is no universal “right answer” for these trials. Such trials are called “non-dominated.” We also include “dominated” trials in the experiment, in which one alternative is an objectively better choice than the others, but those serve mainly as catch trials, to assess whether participants are attempting to perform the task to the best of their abilities.

The first main goal of this study is to investigate how different model classes perform when the complexity of the task varies. Task complexity is lower for the 2×2 task than for the 4×4 task, see Figure 1a. This is reflected in the dramatically different behavioral measures of human behaviors, specifically the numbers of saccades and reaction times, both of which increase significantly from the simple to the complex task, see Table 1. Thus, the first question we ask is how the predictive power of different model classes varies for varying task complexity.

**Table 1:** Median number of fixations and mean reaction times (RTs) for tasks of different complexity. Only data from non-dominated trials are included.

| Task type | Number of Fixations | RT [ms] |
| --- | --- | --- |
| 2opt-2att (2x2) | 5.6 | 3,541 |
| 4opt-4att (4x4) | 20.7 | 14,214 |

Of the different computational models we develop in the present study, some do make use of the eye movement observations while others do not. The second main objective of this study is to determine to what extent information about the (overt) attentional state of individual participants in individual trials improves model performance in the prediction of this individual’s eventual choice at the end of the trial. We will also look at the combination of the two main questions, *i.e.* whether there is an interaction between complexity of the task and availability of attentional data for the predictive performance. For instance, does knowledge of eye movements improve model performance when task complexity is high compared when it is low?

### 2.2 Overview of computational models

Our goal is to understand decision making in tasks of varying complexity (Section 4.3) by developing quantitative models of the underlying processes. We here provide a brief overview of all models considered before describing them in detail in Sections 2.3-2.10. In Section 2.3, we consider baseline models that are determined only by the overall choice behavior, *i.e.* the choices participants make over all trials (or over all trials in the training set, used to determine model parameters which are then tested on the test set). The most simple concept is maximizing the expected value (EV) of collected rewards (amounts). A simple generalizations of this is maximization of the Subjective Value (SV) which is computed as the sum of weighted attribute values for a given alternative; we call this the Additive Rule (AR). Alternatively, the SV can also be computed as a nonlinear combination of attribute values. This approach is taken in Prospect Theory (PT). In its baseline version, this model only uses choice behavior without making use of eye movement data. In other models discussed below, SV will be computed based on choice behavior and then used on a fixation-by-fixation basis.

A model that takes into account behavior in more detail is studied in Section 2.4. Rather than determining behavioral strategy from the set of all (training) trials, the assumption is that behavior is dynamically influenced by the different outcomes of individual trials. This is represented in the formal model by a number of internal latent variables related to the reward history that contribute to the computation of the subjective value. Optimizing this value then introduces dynamic, history-dependent changes in behavior.

Subsequently, even more fine-grained influences are included, at the level of eye fixations rather than trials. In Section 2.5, we assume that participants perform Bayesian inference computations based on the set of fixations to optimize either EV or SV. Finally, we introduce model classes where not only the set of fixations can influence choice behavior but also their sequence. In Section 2.6, models in the first class use evidence accumulation at each fixation through a drift-diffusion model that is updated at each updated shift of overt attention, *i.e.* at each fixation. Section 2.7 describes Decision by Sampling, a heuristic model of decision making based on a series of binary comparisons between attributes, with the alternative chosen that has the highest number of favorable comparisons. In contrast, the Leaky Accumulator model, discussed in section 2.8, assigns values to each available alternative and finally selects the highest-valued. Integration of value occurs by attentional selection of attributes and it is a lossy process. The final two models are inspired by work in .Operations Research. Section 2.9 is an implementation of the Preference Ranking Organization Method for Enrichment Evaluations (PROMETHEE) algorithm. We modify this model by taking into account the sequence of information selection determined by eye movements as well as memory leaks to arrive at truncated-normal PROMETHEE model (TNPRO) in Section 2.10.

Predictions of all models were tested on data collected in the tasks described in Section 4.3. Briefly, in “2×2 experiments”, participants selected one of two alternatives where each alternative was characterized by two attributes, the probability *p* to obtain a reward of size *x*. In “4×4” experiments, a choice was made between a total of four alternatives, with each having a probability *P* to obtain a reward of size *A* (as before) and, in addition, the probability of losing an amount *L* with probability 1 − *P*, and having to wait for a delay *d* until the outcome of the gamble is revealed. For all models, the task was to predict the set of alternatives chosen by each participant.

### 2.3 Baseline models

#### Expected Value

By elementary probability theory, the expected value of the winning amount for each alternative in the 2×2 experiment is EV = *x* · *p*. In the 4×4 experiment, the expected value to win is

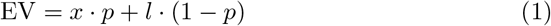

In our experiment, participants have to wait for a delay time *d* before the result is revealed to them. We assume that they experience a simple linear discounting factor which is modeled by subtracting a term proportional to the waiting time, thus

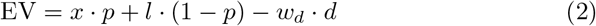

with *w_d_* a free, participant-specific parameter. The prediction is that the alternative with the highest EV is selected by the participant. The same applies to the subjective values, eqs. 3 and 6, and for AR, eq 7.

#### Subjective Value

The subjective value in the 2×2 experiment is defined from prospect theory [18, 43] as

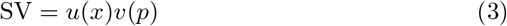

where *u* and *v* are the utility function and weighting function for probability, respectively. Their definitions are:

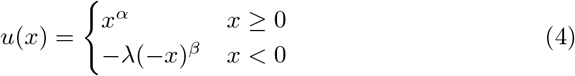

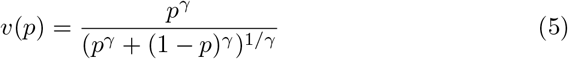

with *α, β, γ, λ* participant-specific parameters. The utility function *u* captures loss aversion. The weighting function *v* is an S-shaped curve that for positive values of *γ* overestimates small probability values and underestimates large probability values. Obviously, for *α* = *β* = *γ* = 1 we have *SV* = *EV*.

In the 4×4 experiment we assume temporal discounting as before. The subjective value is then a combination of the products of utility *u*(*x*) and weighted probability *v*(*p*) and the linear discounting function,

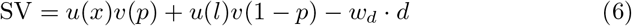

#### Weighted addition: The AR model

In the two models discussed so far combine amounts and probabilities multiplicatively. In our last baseline model, we assume additive combinations instead. For the 4 × 4 case, the quantity to maximize is then the linear combination of attributes

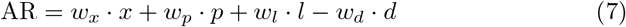

with all coefficients *w* being weighting parameters and *w_l_* = *w_d_* = 0 for the 2 × 2 experiment. [27] called this the “weighted additive (WADD) rule,” we will simply call this the additive rule, AR. Although not maximizing the expected amount to win, this is a simple heuristic which may be used by participants. In a separate experiment, not discussed further here, we found that this model has high predictive power for the behavior of monkeys performing a similar task.

In the subsequent descriptions of additional models, Sections 2.3-2.10, we will describe models mainly for the 2×2 experiment and not always include explicitly the additional parameters for the 4 × 4 experiment, *d* and *l*. In all cases, we will assume the same linear temporal discounting function as in this section.

### 2.4 Intertrial effects modeled by latent variables

This model maximizes SV, as in section 2.3, but it allows that participants may dynamically change their strategy based on the feedback given at the end of each trial (see section 4.3). One example of such influences is the “hot-hand fallacy,” in which a participants experiences a “string of lucky events” and takes more risks than in a more neutral situation. The opposite is the “gambler’s fallacy” where a participant’s behavior is guided by the feeling that there is a limited number of positive outcomes, that this number may be (temporarily) exhausted and, as a consequence, makes less risky choices. In our case the feedback is the money reward *R* announced after the time delay *d* (note that *R* ∈ {*x, l,* 0} due to the stochastic nature of our task). The change of the decision making strategy is modeled by updating the parameters of the utility function *u* and the probability weighting function *v* after each trial. For example, increased risk-seeking increases *α* and decreases *γ* in eqs. 4 and 5. The influence over the trial history, with more recent trials weighted higher than more temporally distant ones, is implemented by means of the latent variable (*LV*) as the running average of the feedback impact,

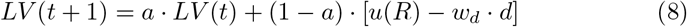

In this equation, *a* ∈ [0, 1] is a decay parameter fitted for each participant. In the extreme case *a* = 0, only the result of the immediately preceding trial has an effect while for *a* approaching unity *LV* will maintain a long-term memory of previous reward history with little influence from the immediately preceding outcomes. The time-discounted utility *u*(*R*) of the reward *R* is *u*(*R*) − *w_d_* · *d*.

All parameters of the temporally discounted SV computation are modified by *LV*. Before the first trial, the parameters of SV are computed based on all trials by fitting all free parameters in eqs 3-5 to best explain observed performance. Then these parameters are considered time-dependent variables that are updated after each trial according to

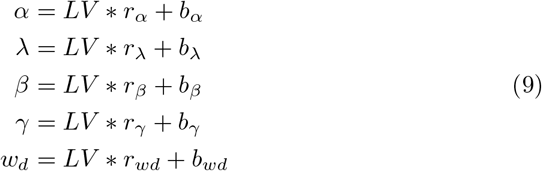

Thus, the parameter values *α* to *w_d_* are linear functions of the latent variable *LV*, each with an intercept *b_x_* and a slope *r_x_*, where *x* ∈ {*α, β, γ, λ, wd*}. In the special case of all variables *r_x_* equal to zero and all variables equal to their intercepts, this model is the original subjective value model. As before, the models predicts that the alternative with the highest SV is selected.

### 2.5 Bayesian inference models

#### Bayesian inference based on EV

There is strong evidence that overt attention plays an important role in the decision process [19, 9]. We take fixations into account in a Bayesian formalism in which the information gained in each fixation is used to update the estimate of the EV. Let *ev* be the expected alternative values obtained from the observed attributes in the fixations of a trial. By Bayes Rule, the estimate of EV is updated by the value *ev* observed in this trial by,

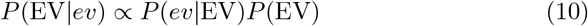

where *P* (*ev*|EV) is the likelihood of observing the fixated values *ev* given the expected value *EV*, and *P* (EV|*ev*) is the posterior distribution of the expectation value given the observed values *ev*. Under the simplifying assumption that fixations are independent, the likelihood is a Gaussian with mean and variance determined by the set *ev*. The *ev* values are computed from the observed (fixated) attributes according to eq. 2. If an attribute value has not yet been observed, it is replaced by the default which is 0.5 for *x*, *p*, and *d*, and −0.5 for *l*. The prior *P* (EV) is the estimate of EV before any data is observed which is computed from the priors of the attributes used to compute the *ev* values. We assume that the attribute priors are all uniform (flat). For the simplest case, the 2 × 2 experiment, the product of the two uniform distributions for *x* and *p* results in a logarithmic distribution for the prior.

#### Bayesian inference based on SV

Bayesian inference can also be performed on SVs which are computed from eqs 4 and 5 over all trials of a participant. Attribute values are then transformed into SVs analogously to the EV case, with the same initial conditions, and Bayes Rule is then

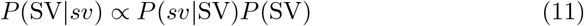

The prior distribution for *SV* is more complicated and computations are all performed numerically. Details are included in the appendix.

### 2.6 Attention-modulated Drift-Diffusion Model (aDDM)

A successful class of cognitive models are race models, or the closely related drift-diffusion models [29] which enjoy neurophysiological support in tasks like perceptual decision making [11]. The basic idea is that the brain gathers evidence over time and when sufficient evidence for one of the alternatives has accumulated, the decision is made to choose this alternative. Our experimental paradigm is particularly suited to test this type of model because the process of evidence accumulation is well-defined and accessible for quantitative characterization: perceptual evidence is only available during fixations, and we can directly observe which information is becoming available during each fixation. Following [19], we assume that the process of evidence accumulation is influenced by the sequence in which information is obtained, as well as by the duration a given piece of information is attended. This means that in the race towards the decision threshold the slope of an attended attribute is increased relative to that of non-attended attributes in proportion to the time that the attended attribute is fixated.

We discretize time *t* in steps of 1 millisecond and evidence for each alternative is assigned to one accumulator. When a fixation to an attribute occurs, the accumulator for the alternative that the fixated attribute belongs to is updated according to the rules below, eqs 12-15. If, *e.g.*, the fixated attribute is the amount *x*, for the duration of the fixation the accumulator value of this alternative is advanced at discretization step (*t* − 1) → *t* by

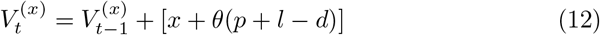

The equations for the other attributes, probability *p*, loss *l*, or delay *d* are analogous,

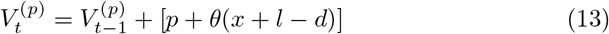

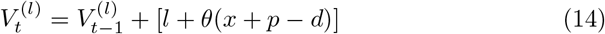

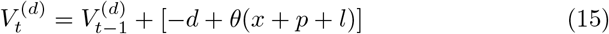

It can be seen that the value of the fixated alternative is increased by the weighted sum of *all* its attribute values. The parameter *θ* ∈ [0, 1], fitted for each participant, is the relative weight by which the contribution of the non-fixated attribute is attenuated relative to the fixated one. For the 2 × 2 experiment, only eqs 12 and 13 apply. In all cases, the alternative with the highest value after all fixations is the predicted choice of this model.

### 2.7 Decision by Sampling model

The Decision by Sampling model (DbS) [35] provides an account for how people make decisions which is entirely based on series of binary comparisons between attributes. The decision between alternative is based on the number of favorable comparisons for each alternative. More specifically, a target attribute is randomly selected from all the attributes in all alternatives. A comparison attribute is randomly selected from either the immediate context (the choices in front of the decider) or from long-term memory. Then a binary, ordinal comparison is made between these two attributes, *i.e.* it is determined which one is better. Subsequently, the value of the accumulator of whichever alternative is more favorable than the other is increased by one. If the difference between accumulator values for each alternative reaches threshold, the alternative with the highest accumulator is selected, otherwise the next target attribute is chosen, as described above, and sampling continues.

#### 2.7.1 Exhaustive Decision by Sampling

Without considering eye movements, we can apply the DbS model directly to our experiment. We assume the sampling ends and the decision is made when an accumulator reaches a fixed threshold, *T*.

While an iterative process as described above could be used, the problem can actually be solved in a closed form in the case of exhaustive sampling, *i.e.* when all possible pairwise comparisons are made. This transforms the model to a simple exercise in permutations. Generalizing the case of two alternatives [35] to our 4 × 4 experiment, the probability for picking alternative 1 is:

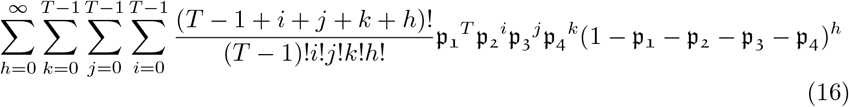

where 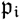 is the fraction of pairwise comparisons in which alternative *i* is favorable over all other alternatives for a given trial. The expression 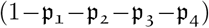 is the probability of no increment, which is the case of an unfavorable comparison with a sample from long-term memory. As noted, 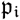 depends on the probability of a favorable comparison to the comparison attribute, which could be sampled from immediate context or long-term memory. In our case, the immediate context consists of the values of the same kind attributes from other alternatives and varies from trial to trial. Long-term memory reflects participants’ previous life experience and, for simplicity, is modeled as a uniform distribution of attribute values. While originally a more complex distribution was used [36], Stewart and Simpson [38] showed that using a uniform distribution has only a slight effect on model predictions, as does choosing different stopping rules. The probability of sampling a comparison attribute within the immediate context, rather than from memory, is a free parameter 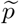.

#### 2.7.2 Decision by sampling with known attentional influence

We can expand the DbS model to the case when eye movement data are available because we then know which target attribute is selected during each trial. Now we only need to sample the comparison attribute randomly, either from immediate context or from long-term memory. To be specific, for every fixation within the trial, we randomly sample a comparison attribute and compare the fixated attribute with a comparison attribute. As before, the probability of sampling from the immediate context is 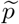. If the comparison is favorable, the accumulator of the fixated alternative will be incremented by one count. At the end of the trial, the model predicts that the alternative with the highest accumulator count is chosen. Note that no stopping rule is needed here because the fixation history is known. To obtain an estimate for the probability of selecting each alternative, this process is repeated *N* times for every trial, with *N* chosen appropriately, *e.g. N* = 200. By counting the number of times the model makes different predictions we obtain the probability for each alternative.

### 2.8 Two-layer leaky accumulator model

Leaky accumulator models successfully explain several behavioral patterns of multi-alternative choice tasks [45, 46]. Glickman *et al.* introduced a leaky accumulator model explaining the formation of preference in risky choice [9]. The model consists of two layers of accumulators where the first collects subjective attribute values and the second integrates the information of each alternative. Under our 4 × 4 experiment setup, the first layer consists of 16 accumulators, 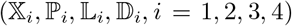, representing the activations of different values (*x_i_, p_i_, l_i_, d_i_, i* = 1, 2, 3, 4). These accumulators are updated according to the following equations:

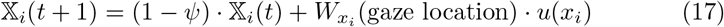

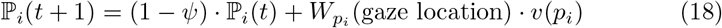

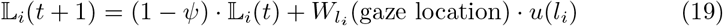

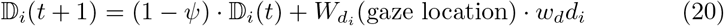

where *t*, a measure of time, denotes the index of fixations from the beginning of the trial. The constant *ψ* ∈ [0, 1] represents degradation of information over time and *u, v, w_d_* · *d* are subjective value functions defined in the Subjective Value model, section 2.3. The step function 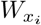 is defined as,

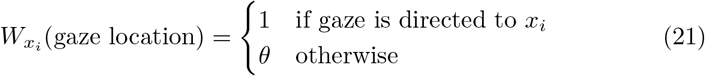

where *θ* ∈ [0, 1] quantifies the decreased influence of attribute *x_i_* when it is *not* attended. The functions 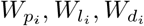 are defined analogously to 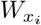.

The second layer consists of four leaky accumulators 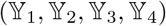 integrating the subjective values of the four alternatives according to the following difference equations:

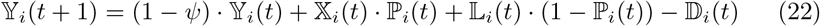

where *ψ* and *t* are the same as in the first layer. All accumulators are initialized to 0 at the beginning of each trial.

### 2.9 The PROMETHEE model

The Preference Ranking Organization METHod for Enrichment Evaluations (PROMETHEE) mode is a class of outranking methods in multicriteria analysis [7], which are used to rank multiple alternatives given many criteria. In this model, a multicriteria preference matrix **M** is first built from the weighted sum of preferences between attributes for pairwise comparisons between alternatives. In our 4 × 4 experiment, the matrix element of **M** in the *i^th^* row and *j^th^* column is computed as,

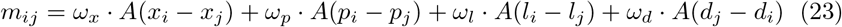

This element represents the strength of the preference the decision maker has for alternative *i* over alternative *j*. The coefficients (*ω_x_, ω_p_, ω_l_, ω_d_*) are the weights for attribute types (*x, p, l, d*). The function *A* is defined as,

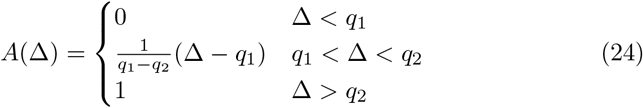

which expresses the result of comparison in terms of preference. There are many other possible forms for this function [7, 6] but we choose this piece-wise linear form for simplicity. The free parameters *q*1, *q*2 define the portion of the function with finite slope and determine how differences in comparison influence the intensity of preference.

By construction, the sum over all elements in row *i* is the summed preference of alternative *i* over all others. Likewise, the sum over the elements of column *i* is the summed preference of all alternatives over alternative *i*. The net “outranking flow” for this alternative is defined as their difference,

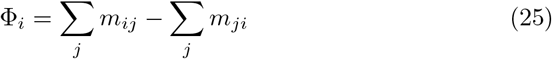

The model predicts that the alternative with the highest net flow is selected.

### 2.10 The TNPRO model

In the original PROMETHEE model, defined in section 2.9, the decision is made based on the available alternatives and attributes. This model does not take into account the *dynamics* of decision making. In this section, we extend the PROMETHEE model to take into account the attentional state while observers gather information and arrive at their decision. The extensions that we implement are (i) adding two leakage components to account for lossy memory contents and (ii) selecting a specific weight function for the attribute sums, namely the fixation duration. We also focus on the special case of using truncated normal distributions for attribute values and memory contents.

Inspired by the PROMETHEE model, we adopt the outranking flow methodology from that model but extend it by including a lossy memory component. We also take advantage of the structure of our decision space in which variables are Gaussians over a finite interval, assumed by us without limit to generality as [0, 1]. Since these Gaussians are probability distributions they are normalized. We call the model therefore the **T**runcated **N**ormalized Gaussian **Prom**ethee, or **TNPRO** model. Furthermore, we take into account the dynamically changing state of overt attention by building up the advantage matrix used in the outranking procedure during each trial by using the information gathered at each fixation.

#### 2.10.1 Memory model

Our model predicts choices based on fixations and attribute values for each trial. The obtained attribute information is not necessarily perfect because of perceptual, processing and storage noise.

Our assumption is that the value retrieved from memory is a Gaussian distributed random variable *X* whose mean *μ* is the true attribute value and that has a time-varying standard deviation *σ*(*t*). The distribution needs to be restricted, however, over a finite range since all values are positive and also have a finite maximum value. Without loss of generality, we assume the latter to be unity. Inside the interval *x* ∈ [0, 1] the distribution is then,

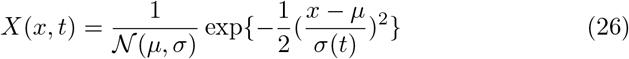

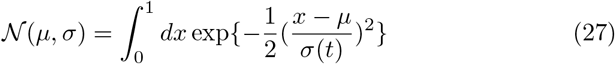

and it is zero outside this interval. A zero value for *σ* would mean certainty of the stored value, and the value’s mean 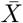 would be identical to the true attribute value.

Before the first fixation of an attribute no information is available and the distribution over the available range of attribute values is flat. At the time of each fixation of attribute *X*, *μ* is updated to the observed value of this attribute and we set *σ* = *σ*_0_ which is a parameter fitted for each participant. Over time, unless this attribute is fixated again, fading memory results in an increase of *σ*(*t*), representing increasing uncertainty of its remembered value. We assume that the rate of forgetting is highest early on and that it decreases over time; specifically that the standard deviation *σ*(*t*) of the Gaussian is inversely proportional to itself,

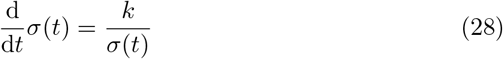

or equivalently,

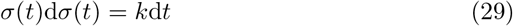

where *k* is another participant-specific free parameter. After integration over both sides of eq 29, the time-dependent width of the distribution is given by

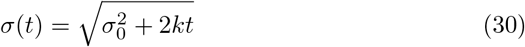

The top panel of Figure 2 shows schematically how contents are read into memory and decay over time. An important consequence of the finite range of the probability distribution is that increasing uncertainty (larger *σ*(*t*)) moves 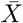 closer to the center of the possible range, resulting in an estimation bias due to forgetting. This is illustrated in Figure 3.

**Figure 2:**
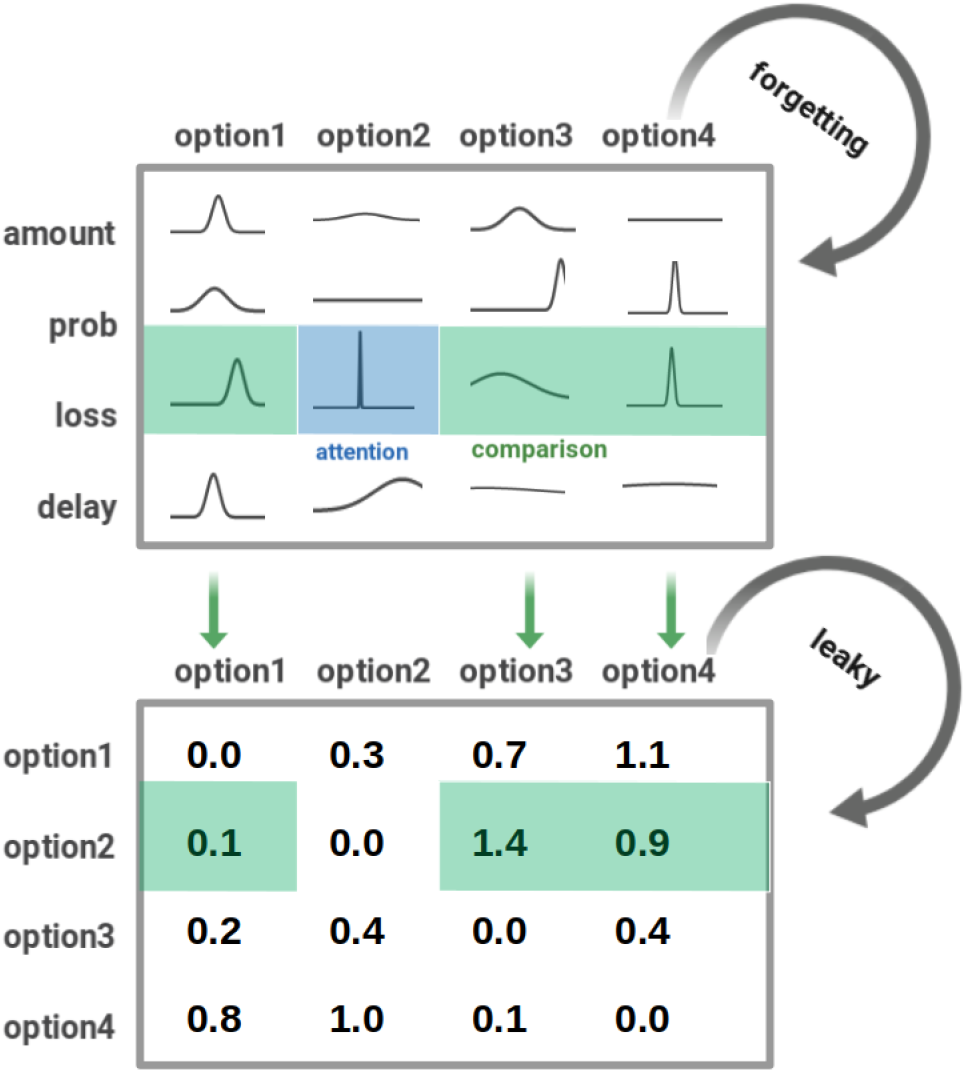
Schematic diagram of information flow in the TNPRO model. The top panel shows the memory model in the case of four alternatives (columns) and four attributes (rows). The distribution of the remembered value is shown for each of the 16 attributes. In the example shown, a fixation (“attention”) has just occurred on “loss” attribute of alternative 2, resulting in a very tight distribution (*σ* = *σ*_0_) of this attribute. The remembered values of other attributes have broadened over time, with their standard deviations given by eq. 30. The value of this attribute is compared against the corresponding attributes of the other alternatives shown in the green shaded areas (“comparison”) and used to compute the elements of the matrix **M** from eq. 23, section 2.10.2. Memorized values are used in this Advantage matrix (bottom panel) which is participant to memory loss itself. See text for details.

**Figure 3:**
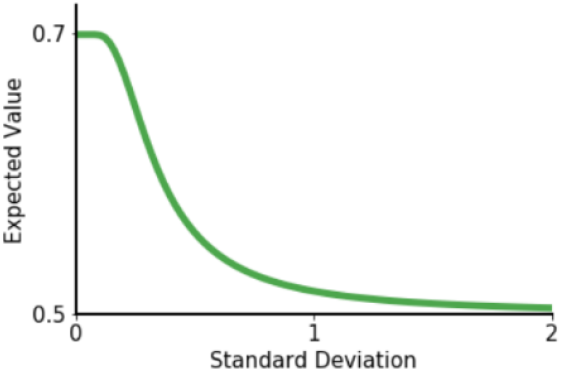
Increased variance of a truncated Gaussian leads to a bias towards the center of its range, *i.e.* 0.5. Shown is the mean of a Gaussian distribution limited to the range [0, 1] and centered at 0.7 as a function of its standard deviation *σ*. Already for *σ* = 1 the mean is very close to 0.5.

**Figure 4:**
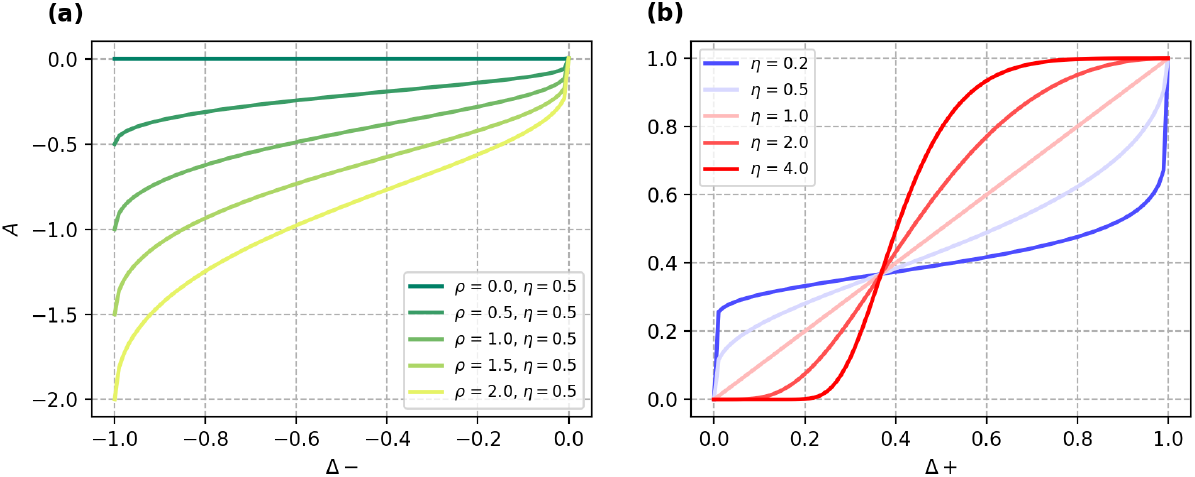
Advantage function, eq. 32, for several values of *η* and *ρ*, plotted separately for negative (a) and positive (b) arguments. This captures the observation that some people are sensitive to small difference while others are not.

**Figure 5:**
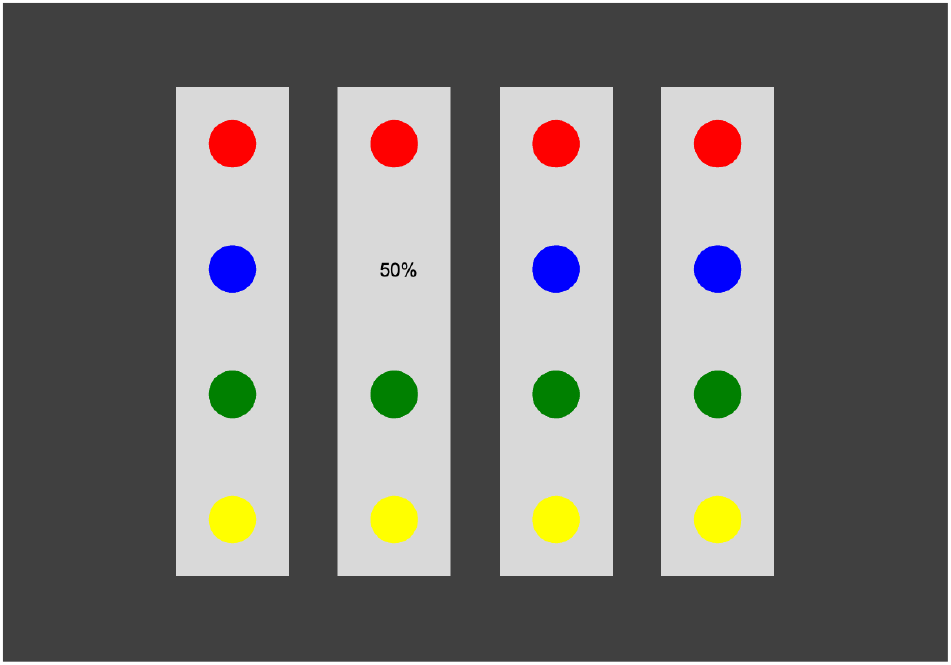
A stimulus of one trial on the 4 alternatives 4 attributes task. Red, blue, green, and yellow indicate loss amount, winning probability, winning amount, reward delay, respectively. The valuie of the winning probability of the second alternative is revealed.

#### 2.10.2 Determination of relative attribute values

At every fixation, the values of all attributes are accessed and updated in memory, see section 2.10.1. The value of the fixated attribute is compared to that of the corresponding attributes belonging to all other alternatives as retrieved from memory. Assume the fixated attribute is of type *π* ∈ {*x, p, l, d*} in alternative *i*. We use *X_π,i_* for its attribute value which is one of the values *x_i_*, *p_i_*, *l_i_* or *d_i_*. The difference Δ_*π,ij*_ between the attended attribute *X_π,i_* and the corresponding attributes *X_π,j_* for another alternative *j* is defined as

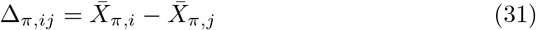

where 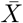 is the expected value of attribute *X*, computed as the mean of the corresponding distributions from the memory model defined in section 2.10.1. The 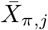 values are retrieved from memory and therefore subject to a decay in accuracy, characterized by the width *σ*(*t*) of their Gaussian spreads, eq. 29.

When the attended (fixated) attribute is better than the corresponding attribute of alternative *j*, *i.e.* Δ_*π,ij*_ > 0, the former has an advantage over the latter. The function to quantify this advantage has a range [0, 1] where 0 means that both attributes are of equal value and 1 means that the attended attribute is preferred in the strongest possible terms. The “Advantage” function we choose is

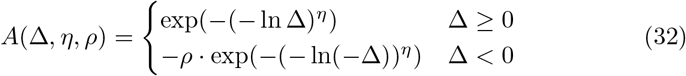

and it is shown in Figure 4.

This function is asymmetrical around the origin because values of *ρ* different from unity differentiate between positive and negative differences between alternatives. If *ρ* ∈ [0, 1] as we usually find (see below), disadvantages of the attended attribute relative to the remembered attribute of other alternatives are weighted lower than their advantages. As a reminder, this asymmetry is extreme in the original PROMETHEE model [6] (see also section 1.3) where the advantage functions are strictly zero for negative arguments, corresponding to *ρ* = 0 in eq 32. We relax this restriction and, instead, fit the parameter *ρ* for each participant. While the best value for this parameter was usually not zero, we did find it that typically takes on relatively small values, see section 3.8. The attended glass seems to be seen as half-full.

#### 2.10.3 Relative alternative values based on differences between attributes

Section 2.10.2 defines comparisons between individual attributes. The completion of the decision process requires, however, that eventually one of the *alternatives* is chosen. In this section, we describe how the relative advantages of different attributes are used to compute relative differences between alternatives.

In the original PROMETHEE model, section 2.9, the integration of attribute differences into alternative values was achieved by the weighted sum in eq 23. In the TNPRO model, relative alternative values are constructed and updated successively during attentional information gathering, *i.e.* based on observed eye movements. As in the PROMETHEE model we organize these into a matrix **M** whose element *m_ij_* indicates the subjective advantage of alternative *i* over alternative *j*. The matrix is illustrated in the bottom panel of Figure 2. We assume that at every fixation the participant compares the fixated alternative with all other alternatives, based on the attribute type *π* (*e.g.* amount to win, probability to win, *etc.*) that is being observed in this fixation. If there are *N* available alternatives, this makes (*N* − 1) pairwise comparisons. For each pairwise comparison, the matrix element *m_ij_* is updated based on the subjective advantage that the fixated attribute (type *π*) of the fixated alternative, *i*, has over that of alternative *j*,

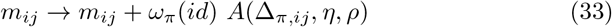

using *A*(Δ_*π,ij*_, *η, ρ*) from eq 32 and Δ_*π,ij*_ from eq 31. The participant- and attribute-dependent weighting factor *ω_π_*(*id*) is defined in the following paragraph.

The relative emphasis put on the different attribute types (*e.g.* risk averse *vs.* risk-seeking) may vary on a participant-to-participant basis, and this emphasis may also change over time (*e.g.* due to response feedback; examples for such history-dependence are the hot-hand fallacy and the gambler’s fallacy [28, 34]). We assume that the importance of attributes is reflected in the amount of attention directed to them, which is in our paradigm the number of times an attribute is being fixated. Explicitly, our assumption is that attributes of lesser importance to a given participant at a given time will be fixated less often than more important ones (see [20] for a similar idea). The contribution of attribute type *π*, *A*(Δ_*π,ij*_, *η, ρ*), is therefore weighted by *ω_π_*(*id*), the total number of fixations spent fixating attributes of type *π* up to the *id^th^* trial.

The final step in the construction of matrix **M** takes into account that newly formed preferences will in general have more impact on the decision than previously formed preferences (recency effect). We therefore include a participant-specific fading parameter *δ* ∈ [0, 1] by which all matrix elements are multiplied after every fixation,

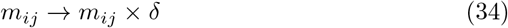

#### 2.10.4 Alternative ranking and choice prediction

We now show how the set of pairwise differences between alternatives captured in **M** is used to obtain a ranking between alternatives. Slightly generalising a concept from the PROMETHEE model, eq 25, we define the participant-specific net outranking flow for optiom *i* as,

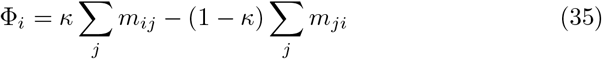

where *κ* ∈ [0, 1] parameterizes the participant-specific difference with which positive and negative outranking flows are weighted. This takes into account that some individuals may focus more on the positive aspects of an alternative while others are more influenced by how much other alternatives are preferable.

The alternative with the highest Φ_*i*_ is predicted to be chosen, *i.e.* max_*i*_(Φ_*i*_).

Our model uses a truncated normal distribution as a model of working memory, and then updates its advantage matrix in a way similar to the PROMETHEE algorithm [7]. Therefore we call our model the “Truncated Normal-PROMETHEE model,” or “TNPRO model.”

Some of the intuitions in our model are similar to those in a context-dependent model by [44] which combines the effect of background context (alternatives encountered in the past) and local context (offered alternative set in current trial). There are also several important differences. First, in their model, the background affects the *global* change in the relative weight of the attributes, while in our model this is captured by *trial-specific* attribute weights *ω_π_*(*id*). Considering there could be more factors contributing to this weight change, such as the hot-hand fallacy and gamblers’ fallacy resulting from the instant feedback at the end of each trial, we chose to use proportion of fixations as a more general approach to capture possible latent variables across trials. A second assumption in their model is that the effect of local context can be interpreted as a “tournament” in which the candidate alternative is matched against all the other presented alternatives, and its overall score is the sum of the results of these matches. This is also the case in our model where several pairwise comparisons occur at every fixation and the final decision is made from the accumulation process of participantive advantage values. In contrast to our model, however, Tversky and Simonson assume that the disadvantage of alternative *i* to alternative *j* should have at least the same impact as the advantage of alternative *j* to alternative *i*. In our model, we are agnostic to the relative impacts and allow them to vary unconstrained between participants (parameter *κ*, eq 35). We actually find the opposite of what is assumed by [44], namely that *ρ* usually gives a much smaller value than unity, which means that the net impact of advantages of an alternative strongly dominates that of its disadvantages. We also implemented a simplified model, with *κ* = 0 in eq 35 and found that its performance was only marginally worse than the full model. Its prediction error was consistently higher than that of the full model (with fitted *κ*) but only by less than 0.5% (data not shown).

### 2.11 Parameter estimation

In the 2 × 2 experiment, we have 540 trials per participant on average and in the 4 × 4 experiment we have 168 trials per participant on average. The participants’ data are divided into a training set and a testing set. For each participant, the first 80% of the trials are taken as training set and the last 20% are taken as test set. The free parameters in the models are optimized for each participant to fit the choice made at every trial in the training set. We determined these parameters separately for each participant by maximizing the likelihood [46] of generating the correct choice prediction. Because the likelihood function typically has many local minima, we used a simulated annealing algorithm, implemented by the *dual annealing* function from the Python scipy.optimize package. The initial “temperature” parameter of the algorithm was set to 15,000, a relatively high value, to allow access to a large part of the energy landscape. The maximum number of global search iterations was set to 2,000 for all the models and the optimization process was terminated if this number of operations was reached. Cross-validated test sets are created by…

AIC and prediction accuracy on the training set are calculated. On the test set, the log likelihood and prediction accuracy are computed. The results are presented in Table 2.

**Table 2:** Summary of results for the 2 × 2 (top) and 4 × 4 (bottom) experiments for all models. Shown are Akaike Information Criterion (AIC), choice prediction accuracy for the training set, negative log-likelihood, and choice prediction accuracy for the test set. AIC values are rounded to the nearest integers. AIC differences exceeding 10 are considered very strong evidence in favor of the model with the lower numerical values. Bold entry indicates the best fitting model for each measure.

|  |  | AIC | Prediction Accuracy | - 2 Log-Likelihood<br>(cross-validated) | Prediction Accuracy<br>(cross-validated) |
| --- | --- | --- | --- | --- | --- |
| $2 \times 2$ | EV | 10729 | 84.0% | 2638 | 84.2% |
|  | SV | 8592 | 89.5% | 2070 | 89.8% |
|  | AR | 9148 | 87.4% | 2115 | 88.4% |
|  | BI-EV | 9869 | 86.8% | 2361 | 87.4% |
|  | BI-SV | 8795 | 88.8% | 2095 | 88.7% |
|  | LatentVariable | 8708 | 89.5% | 2047 | 89.6% |
|  | aDDM | 9634 | 87.0% | 2365 | 87.4% |
|  | DbS | 10699 | 81.4% | 2549 | 82.1% |
|  | DbS with attention | 9730 | 85.7% | 2391 | 86.1% |
|  | **Two-layer leaky accumulator** | **6435** | **92.7%** | **1614** | **91.8%** |
|  | PROMETHEE | 8735 | 87.4% | 2025 | 88.2% |
|  | TNPRO | 8392 | 88.9% | 2074 | 88.6% |
| $4 \times 4$ | EV | 2646 | 78.3% | 694 | 73.9% |
|  | SV | 2058 | 85.2% | 594 | 81.7% |
|  | AR | 2113 | 84.0% | 648 | 78.7% |
|  | BI-EV | 3526 | 76.6% | 902 | 74.6% |
|  | BI-SV | 3683 | 70.2% | 868 | 77.1% |
|  | LatentVariable | 2132 | 86.4% | 658 | 79.0% |
|  | aDDM | 4173 | 65.1% | 978 | 71.9% |
|  | DbS | 4922 | 37.5% | 1626 | 40.0% |
|  | DbS with attention | 2526 | 78.4% | 643 | 76.7% |
|  | Two-layer leaky accumulator | 2304 | 88.9% | **389** | 86.5% |
|  | PROMETHEE | 2147 | 84.6% | 754 | 79.6% |
|  | **TNPRO** | **1648** | **89.1%** | 461 | **87.0%** |

We compare the performance of our models and we analyze the parameter values fitted to the experimental. Since parameter values are not necessarily distributed normally, non-parametric tests are used to find statistical patterns, at the cost of some statistical power. Spearman rank-order correlation coefficients, *r_s_*, are calculated to quantify correlations between parameters. Wilcoxon rank-sum tests are used to assess the difference between parameter values from Expt. 2 × 2 and Expt. 4 × 4. The following conclusions are our interpretation based on the statistical results.

## 3 DISCUSSION

In this report, we study the performance of 12 model classes for predicting choices of human participants in two sets of gambles of different complexity. For the set consisting of simple gambles (2 alternatives with 2 attributes each), we find a relatively narrow range for the prediction accuracy. On the training set, prediction accuracy varies from a low of 84.2% (for EV) to a high of 91.8% (for the leaky accumulator). The range is 81.4% (exhaustive DbS) to 92.7% (leaky accumulator) for the test set, see Table 2. Though there is some variation, all of these very different model classes seem to be able to predict behavior reasonably well, all making correct predictions for better than eighty, and sometimes ninety, percent of choices. We are possibly looking at a ceiling effect in prediction accuracy that does not allow us to meaningfully differentiate between the models tested. We note that this is not the case for either the AIC measure nor the negative log-likelihood: in both measures, the spread between models is much larger. However, the two lowest-performing models for prediction accuracy also have the worst measures for AIC and neg. log-likelihood, and the best-performing model for prediction accuracy has the best results for these measures.

For the complex gambles (4 alternatives with 4 attributes each), the first observation is that overall prediction accuracy is generally lower. Each model performs less well in this situation than the equivalent model in the simple situation. Overall lower prediction accuracy for a more complex situation is, of course, not unexpected. More interesting is that for the more complex choices, the *differences* between models vary considerably more than in the simpler task. Prediction accuracy on the training set varies from a low of 37.5% (exhaustive DbS) to a high of 89.1% (TNPRO), *i.e.* by considerably more than a factor of two. Results are similar for the test set.

It thus appears that the simple 2-alternative 2-attribute choice, variations of which are used in a large number of studies, may not be very suitable to differentiate between models, at least if choice prediction is chosen as the criterion to distinguish between models. There are at least two possible explanations for this result. One is that all of the different cognitive mechanisms underlying each of the 12 models tested are suitable for solving this task in their specific way, that different participans are using them with comparable success rates, and that therefore all models achieve comparable (and uniformly high) prediction success. The other is the mentioned ceiling effect: the task may be easy enough that it can be solved rather well by all of the models, even though the mechanism each model underlying is a poor approximation of the actual cognitive processes executed by our participants.

In contrast, the pronounced differences of choice prediction performance for the complex task allow us to differentiate between the model classes more clearly. While still simple compared to many real-world decision making problems, this task is much more cognitively demanding, as is reflected in the substantially larger reaction times and number of saccades made by participants, see Table 1.

In the following section, we first determine which models perform best in the two tasks. After that, we analyze which features are likely to contribute to the success, or lack of it, of different models and discuss features that guide the decision process.

### 3.1 The best performing models

For both of the tasks (2 × 2 and 4 × 4), there is one clear “winner,” *i.e.* one model whose performance is clearly better than all other models. In the 2 × 2 task, the Two-layer leaky accumulator exceeds the performance of all other models in all four criteria: it has the lowest AIC score, the lowest negative log-likelihood, and the highest prediction performance on both the training and the test set. Even in view of the possible limitations that this task may have on the value of comparative evaluations of different model classes, as discussed above, this consistency increases the confidence that this model really is superior over the other tested models. This result is also in agreement with a prior finding where the performance of this model exceeded that of all other models in a very similar task on a non-overlapping set of observers [9]. We also note that there is no clear overall “second-best” model, the runners-up are different in all categories: The second-best performing model for test prediction accuracy is SV, for training prediction accuracy it is SV and Latent variable (tied), for negative log-likelihood PROMETHEE, and for AIC it is TNPRO.

We also can identify the best-performing model in the 4 × 4 experiment. The TNPRO received the best scores in 3 categories, and second-best in the fourth. As discussed previously, in this more complex situation the differences between models are much starker than for the 2 × 2 experiment. In particular in terms of prediction accuracy, both test and training, the TNPRO model’s performance was more than twice as high than that of the lowest performing model (DbS). Different from the 2 × 2 experiment, for this more complex task there is a clear second-best model: the Leaky Computing Accumulator model showed second-best performance in both of the prediction rankings, and best in negative log-likelihood.

While not coming out at the top in any category, Prospect Theory is among the best performers. This warrants a comparison of its defining features with those of the TNPRO model. An important difference is that the latter does not make any of the assumptions as Prospect theory, like the specific forms for the computation of expected utility or the nonlinear form of the influence of probability to win. Of course it makes other assumptions, like the specific working memory model we use or the computation of relative advantages. These concepts are, however, closer to being interpretable in terms of neuronal processing than the purely functional constructs of Prospect Theory. We also point out that computation of expected value or utility, of any form, are nowhere required in the TNPRO model. It has been argued many times that this is a difficult quantity to assign to individual alternatives while value *differences*, which are fundamental to the PROMETHEE and TNPRO models, are much easier to determine. Possibly, for the simple task it is possible for at least a sizable fraction of participants to compute some approximation of an explicitly alternative value (EV, utility, *etc.*) resulting in good results for models based on this, like leaky accumulators or Prospect Theory. This may not be possible any more for the more complex case where attribute differences are much easier to compute, favoring the TNPRO model.

The success of the TNPRO model, developed from the original PROMETHEE model by adding components inspired by biological information processing, like working memory, highlights the value of models that were pragmatically designed in management science to solve complex problems.

### 3.2 History of attentional selection strongly affects decisions

One of the main questions we want to address in this project is whether the detailed sequence of eye movements, *i.e.* attentional deployment, does influence human behavior. Alternatively, eye movements could be a random process for gathering information where the order in which any particular piece of information is acquired does not matter. A useful tool to answer this question is a comparison between the two Decision by Sampling models because they are identical, except one takes into account the specific eye track and the other does not. We find (Table 1) that the DbS model with attentional influence performs far better than its exhaustive version where no attention history data of participants are used. The former’s accuracy on test dataset exceeds the latter by 4 percentage points in the 2 × 2 experiment, and by 36.7 percentage points in the 4 × 4 experiment. It is clear that the attentional history contains crucial information for predicting the participants’ decisions, and this is especially important when the task is more complex. A corollary is that participants do not necessarily make rational choices in these tasks, since a rational decision should only depend on the attribute information and be independent of the sequence in which it was acquired.

### 3.3 Weak correlation between trials

We were also interested to know whether the winning/losing history of previous trials has an impact on the strategy deployed in the current trial. This could be due to reasons like hot-hand fallacy or gambler’s fallacy [34]. We therefore developed in section 2.4 a variation of Prospect Theory in which the reward history modifies all model parameters taking into account interdependencies between trial outcomes, the latent variable model.

We found little evidence to support effects of inter-trial correlations. In fact, Prospect Theory with latent variables performs worse than the original Prospect Theory, with no interaction between trials. Not only does the additional free parameters increase the AIC value, but the prediction accuracy on the test set slightly decreases, by 0.2 percentage points for the 2 × 2 experiment and by 2.7 points for the 4 × 4 Experiment. In other words, including trial outcome information from previous trials decreases prediction accuracy rather than increasing it.

Note that, in theory, the latent variable model from section 2.4 is a generalization of Prospect Theory, from section 2.3, and a perfect optimization procedure should reduce the former to the latter if addition of between-trials interactions reduces the models predictive performance. However, this would require that all ten free parameters in eq 9 are set exactly to their required values (all *r_x_* = 0 and all *b_x_* equal to *x*, for *x* ∈ {*α, β, γ, λ, wd*}), in which case the parameter *a* would become irrelevant. We surmise that our minimization procedure is not capable of reaching this global minimum in the high-dimensional landscape of this optimization problem.

Instead, we find that the distribution of the weighting parameter of intertrial interactions *a* becomes bimodal. Its value takes on values either close to unity or close to zero, see section C.6. In the 2 × 2 experiment, 24 of the 34 participants have *a* close to unity, with an average value of 0.998 while for the other ten participants the average is 0.0658, *i.e.* close to zero. A similar pattern can be observed in the 4 × 4 experiment where for 13 out of 16 participants *a* is close to unity. If *a* = 1, the outcome of a trial is not influenced by the outcome (reward) of any preceding trial while for *a* = 0 the influence is limited to the non-weighted sum of all previous rewards, see eq. 8. We conclude that for most people the feedback from previous trials has basically no effect on the participants’ current decision making strategy.

### 3.4 Fixation duration is not a good predictor for decision choice

The aDDM model, defined in section 2.6, assumes that the longer a participant attends a piece of information, the stronger its effect will be on the decision. We do not find strong evidence supporting this hypothesis in our experiment. Judging by the prediction accuracy for the test set, the aDDM model ranks 8th out of 12 in the 2 × 2 experiment and 11th in the 4 × 4 experiment. For the training set, only 65.1% of accuracy on the training set is achieved for the 4 × 4 experiment.

The aDDM model can be seen as a variation of the AR model (section 2.3) in which fixation duration is used to weigh parameters for different attributes. However, the AR model’s accuracy exceeds aDDM by one percentage point on the test set in the 2 × 2 experiment and by 6.8 percentage points in the 4 × 4 experiment, despite just being a simple linear model. Instead of using fixation durations, in the AR model the weight of each attribute is taken as a free parameter. These parameters thus reflect the importance that participants attach to different attributes, at least within the framework of this simple model. The comparison between the two models suggests that fixation duration is not among the factors that determine the decision. A possible explanation is that participants make short but frequent fixations to attributes that are important for their decision. This suggests an alternative model, not explored in this study, in which the number of fixation of an attribute, rather than the total time of fixation of an attribute, is taken into consideration.

### 3.5 Importance assigned to different attributes

The AR model also allows us to determine, within the confines of that model, which attribute participants attach most importance to. For the 2 × 2 experiment, we find the relative means over participants 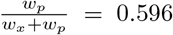 and 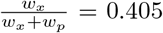. The difference is significant, *p* = 1.95 × 10^−8^ (Wilcoxon rank sum test). Participants thus weigh probability higher than reward. Similarly, for the 4 × 4 experiment, we find 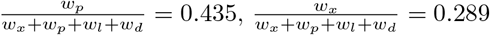, 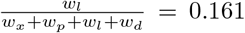, and 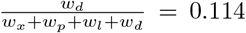. All three differences are significant, with p values of 3.79 × 10^−5^, 1.44 × 10^−5^, 3.00 × 10^−2^, respectively. Thus, the most important factors are probability and reward, in this order, identical to the 2 × 2 case, with loss and delay of subordinate importance.

The analysis above points out that probability attribute plays the most important role in the participants’ decision. Loss amount is given little attention in the 4 × 4 experiment, which is interesting because this shows that an interpretation of the high importance assigned to probability as indication that participants are risk-averse is too simple. This observation is also not easily reconciled with the observation of loss aversion (higher weighting of losses than numerically equivalent wins) in many contexts [18, 43]. It is possible that this effect is due to the details of our experimental design. The probability *p* displayed is that of winning, with the probability of losing being its complement (1 − *p*) which is not displayed. Explicit display of *p*, but not of (1 − *p*) could make winning the more salient outcome. This may suppress the expectation of a loss and have participants focus on the win outcome instead of the size of the penalty in the case of a loss.

Another interesting question is whether the importance given to one attribute is correlated with that assigned to other attributes. By computing the correlation coefficient *r_s_* of the weights on the 4 × 4 experiment, we find negative correlation between 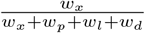 and 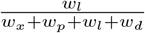, with *r_s_* = −0.688 which is significant, *p* = 3.20 × 10^−3^ (Spearman’s rank order correlation test). Also, 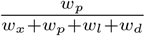 and 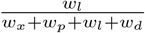 are negatively correlated, with *r_s_* = −0.506 which is also significant, *p* = 4.56 × 10^−2^. Thus, participants who weigh the potential win amount or the win probability highly are less interested in the amount of potential losses, and *vice-versa*.

This means someone would care less about loss if they is interested in either gain or winning probability. This fits our previous explanation about why loss is given little importance. This interpretation is also supported by he parameter values from the PROMETHEE model. In the 4 × 4 experiment, 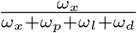 and 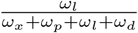 have a correlation coefficient *r_s_* = −0.553, *p* = 2.63 × 10^−2^, and 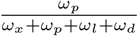 and 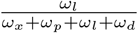 have a correlation coefficient *r_s_* = −0.638, *p* = 7.80 × 10^−3^. Thus, both models show negative correlation between loss and gain/probability. However, there’s not enough evidence showing any correlation between gain and probability in either model.

### 3.6 Impact of non-fixated attributes on choice

Choosing an alternative requires to integrate information from all its attributes, both the instantaneously attended and all others. At any given time, does the attended (fixated) attribute weigh more heavily than the non-attended ones in making the choice, and if yes, by how much? We can get some insight into this question by considering the parameter Θ in the aDDM model, described in section 2.6. This parameter describes the relative impact in the choice of non-attended to attended attributes on, see eqs 12-eqs 15.

We find Θ = 0.515 in the 4 × 4 experiment and Θ = 0.227 in the 2 × 2 experiment. The difference is significant, *p* = 8.78 × 10^−3^, so the attended attributes plays a larger role in the more complex task. One explanation for this difference could be that for every alternative, there are more unattended attributes in the 4 × 4 experiment than in the 2 × 2 experiment (three instead of one), and that the decrease in Θ in the more complex case is needed simply because more components are added. This is unlikely to be the case, however, because the *l* and −*d* terms in eqs 12−15 are negative, within the range (−1, 0). Thus the sum that Θ is multiplied by is not necessarily larger (and could be smaller) in the 4 × 4 experiment than in the 2 × 2 experiment.

Instead, we postulate that Θ goes down as task complexity goes up indicates that in difficult tasks, observers focus on the attended attributes more than on the non-fixated attributes. Given that the latter are not accessible in perception but need to be retrieved from memory, this may be a strategy to economize on working memory under higher load.

### 3.7 The influence of long term memory decays with increasing task complexity

Another interesting question would be: How does a participant evaluate an attribute when looking at it? Is he comparing the attribute with other alternatives, or is he using his life experience to evaluate the attribute?

This problem can be restated using the term in DbS model: Do people select comparison attribute from immediate context or long-term memory? The parameter 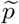 means the probability that participant would sample comparison attribute from the immediate context. For exhaustive DbS model, 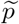 in the 4 × 4 experiment is bigger than the 2 × 2 experiment, with p value=3.77 × 10^−4^. Its mean values are 0.316 and 0.723, respectively. Similarly for DbS model with attentional influence, the mean value for 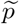 increases from 0.457 to 0.526. This means people are more likely to evaluate attributes using immediate information when task complexity raises. Although long-term memory could help participant to easily evaluate the attribute, it’s more efficient to compare alternatives in immediate context because the goal is to find the best one among given choices. It’s highly probable that the influence of long-term memory would decay as task complexity keeps increasing.

If we combine subsection 3.5 and 3.6, we would reach this conclusion: facing complex tasks, people would tend to only evaluate the fixated attribute, by comparing it with other alternatives in the immediate context. This is exactly what TNPRO assumes what participants are doing. This might explains why TNPRO works so well on the 4 × 4 experiment.

### 3.8 More focus is given to advantages rather than disadvantages

In section 3.6 we analyzed the relative impact of fixated *vs.* non-fixated attributes. We found that for increasing task complexity, the relative weight of non-fixated attributes on the decision decreased significantly. In section 3.5 our analysis of the AR model showed the somewhat surprising result that our participants show little evidence of loss aversion.

We can obtain an alternative view of both effects by analyzing the results of the TNPRO model. The decision process in the first stages of this model is based on pairwise comparisons between attributes. Of particular interest is the advantage that the fixated attribute of the fixated alternative has over the corresponding attributes in non-fixated alternatives. This is quantified in the Advantage function, eq. 32. The parameter *ρ* in its definition determines how much weight is attributed to a situation depending on whether the fixated attribute is better or worse than the corresponding attribute of a non-fixated alternative. They have equivalent weight for *ρ* = 1 and the non-fixated attributes carries less weight for *ρ* < 1. We find *ρ* = 0.327 in the 2 × 2 experiment and *ρ* = 0.0536 in the 4 × 4 experiment.

The fact that *ρ* is smaller than unity in both cases shows that the value of a fixated attribute contributes more to the choice of an alternative when it is favorable compared to when it is non-favorable. This may be the expression of an “optimistic filtering” strategy in which participants aim to find an acceptable alternative, and do not keep detailed track of less-favorable alternatives. This strategy may be needed to decrease the cognitive load by lowering the number of items that need to be attended and/or kept in memory.

This is supported by the fact that *ρ* is much smaller in the more complex 4×4 case than in the 2 × 2 experiment. Dealing with the more complex problems, participants may have to economize cognitive resources more than in simpler situations and therefore jettison information about disadvantages of the fixated alternative more readily. In fact, *ρ* is close to zero for the 4 × 4 case which means that fixation of a non-favorable attribute contributes close to nothing to the decision. It is notable that in the PROMETHEE model, *ρ* is set strictly to zero. We may speculate that this is built into that older model because it is designed to deal with even more complex problems than we do in our experiments.

### 3.9 Small differences are overestimated

The second parameter in the Advantage function of the TNPRo model is *η* which determines the shape of the function, eq. 32. Without loss of generality, we discuss the positive part of the function since the negative part is obtained by mirroring and stretching the positive part. For *η* = 1 the Advantage function is linear. When *η* < 1, the advantage function is convex for small Δ and concave for large Δ while for *η* > 1, the function is concave for small Δ and convex for large Δ. We found that for the 2 × 2 experiment, only 1 out of 34 participants has *η* > 1 and this is the case for 2 out of 16 in the4 × 4 experiment. Thus, most participants overestimate small differences and underestimate the high difference between attribute values.

It is interesting that a function with this general shape is also used for the computation of subjective probability in Prospect Theory, eq. 5. It is not clear whether this is a coincidence since in the TNPRO model, overestimation of small differences applies (on average) to all attributes and not only to probabilities since *η* is a common parameter. Furthermore, in Prospect Theory it is the *value* of the (probability) function that is transformed in this way while in TNPRO the transformation is applied to a *difference* of values.

### 3.10 Retained memory between fixations

A common feature of the two highest-performing models, the two-layer accumulators in the 2 × 2 experiment and TNPRO in the 4 × 4 experiment, is a memory mechanism between fixations. In both cases, memory contents decays exponentially with a tunable decay parameter, *ψ* for the accumulator model and *δ* for the TNPRO model. Note that there is additional memory decay in the TNPRO model (section 2.10.1) which is not taken into account here.

For the two-layer accumulator model, (1 − *ψ*) is a measure of the influence from the previous fixation to the next. Its mean value is 0.268 in the 2 × 2 experiment and 0.409 in the 4×4 experiment. For the TNPRO model, *δ* controls the proportion of the previous advantage matrix that is carried over to the next fixation, see eq 34. Its mean value is nearly identical in both experiments, 0.749 in the 2 × 2 experiment and 0.750 in the 4 × 4 experiment. This shows that the average information retained between fixations is modest. Even for the TNPRO model where the decay is weaker, and for the smallest number of fixations until choice, the influence from the first fixation on the last is less than 0.2 (0.75^5.6^, see Table 1). It is negligible (< 10^−3^) for all other cases.

One possible interpretation is that the decision is highly dependent on the last several fixations. But what is then the use of the fixations early on in each trial? Is the information collected in these fixations essentially not used, and overwritten by the input gathered in later fixations? While we cannot exclude this possibility given the data presented, there is another possible explanation. During the first fixations, participants have no information about the value of many attributes because they have never seen them. Only after a minimum of four (in the 2 × 2 experiment) or 16 (in the 4 × 4 experiment) fixations have been executed can all values potentially be known. We therefore assume that a substantial fraction of participants sample the display using either a random order (because they know the order does not matter) or an idiosyncratic, stereotypical order (because it is easiest to follow the same order in each fixation). The methods applied in the present study do not address this question but we found strong evidence for stereotypical behavior for the first fixations when we performed formal analyses of fixation orders (Elsey *et al.*, in preparation). Since neither these idiosyncratic sequences nor random sequences are correlated with attribute values, the correlation with later fixations (which are presumably chosen based on those attribute values) and with eventual choice are weak at best. This is reflected in the low values for *ψ* and *δ*, respectively, in the two models.

## 4 METHODS

### 4.1 Participants

Participants from the Johns Hopkins University community were recruited to partake in the experiments. All participants reported having normal or corrected-to-normal vision and were able to complete the eye tracker calibration procedure in the 2 × 2 experiment, (n = 34) and the 4 × 4 experiment (n = 24). Written and informed consent was obtained from all participants, and all experimental procedures were approved by the Homewood Institutional Review Board of Johns Hopkins University. Participants were compensated with SONA credit at a rate of 1/hour and given the opportunity to earn a monetary bonus resulting from the outcome of randomly selected trials. Performance >85% on catch trials where one alternative was superior the others was conditional for participant exclusion. In experiment 1, data from 3 participants were excluded from analyses due to performance on catch trials (<85%). In experiment 3, data from 2 participants were excluded from analyses due to performance on catch trials (¡85%).

### 4.2 Apparatus and Stimuli

Participants sat with their head stabilized in a chinrest while eye movement responses were collected using a desk-mounted EyeLink 1000 infrared eye-tracking system (SR Research, Mississauga, Ontario, Canada) at a sampling rate of 500Hz. Participants viewed task stimuli on an Asus LED HD monitor with a 60 Hz frame rate from a distance of 56cm. The behavioral tasks were controlled using a Mac computer (Apple, Cupertino, California, USA) equipped with MATLAB software (https://www.mathworks.com/) and PsychToolbox-3 extensions (http://psychtoolbox.org/).

Stimuli were comprised of several grey bars on a dark grey background. Each grey bar indicated one alternative, comprising of several colored circular masks indicating the attributes. Attribute masks were only extinguished to reveal attribute magnitude when participants were actively fixating upon the respective attribute cue, providing a measure of attention. The alternative bar orientation and attribute masks are rearranged at every trial to control for a potential fixed attribute inspection pattern.

### 4.3 Experimental Task

#### Experiment 1: Two-Alternative, Two-Attribute

Participants performed a risky two-alternative multi-attribute decision making task. Participants were instructed to maintain eye fixation on a central cross to initiate each trial. After fixating for 500ms, the central fixation cross extinguished and two alternatives would appear. Each alternative consisted of two attributes, amount to win and probability to win. Attribute magnitudes were parametrically indicated by visual stimuli in spatially separate locations within a light grey boundary cue indicating that the attributes belonged to the same alternative. Attribute cues were masked by a colored circle (yellow for amount to win; blue for probability to win), masks were only extinguished when participants were actively fixating upon the respective attribute cue. Participants were allowed to inspect the alternatives freely with no time constraint before they indicated their choice with an arrow key press corresponding to the direction of the preferred alternative. Feedback was then provided onscreen as to whether the participant won or lost their gamble.

Alternatives were presented in either combination of three spatial locations (leftward, rightward, upward) lying along an equilateral hexagon (**Figure 4a**). Alternative and attribute locations were randomized on a trial-by-trial basis, resulting in 24 possible spatial configurations. There were 5 different win amount and win probability attribute magnitudes (win amount: .10, .30, .50, .70, .90, probability win: 10%, 30%, 50%, 70%, 90%). 280 trials total were presented over the course of 3 separate blocks. In total, 56 dominated trials (amt1 > amt2 & prob1 > prob2 or amt1 > amt2 & prob1 > prob2) and 224 non-dominated trials (amt1 > amt2 & prob1 < prob2 or amt1 < amt2 & prob1 > prob2) were presented. 64 of the 224 non-dominated trials consisted of alternative pairings with equal expected value (|*EV*| = 0). Overall, the experiment comprised of 19 distinct alternatives sampled from a two-dimensional decision space. A related task, however without eye-movement controlled masking, was described by [9].

#### Experiment 2: Four-Alternative, Four-Attribute

Participants performed a multi-attribute decision making task similar to the 2 × 2 experiment with four alternatives comprised of four attributes. In the 4 × 4 experiment, the two additional attributes were the amount the participant can lose and the delay to feedback of the trial outcome. Attribute cues were masked by yellow (win amount), blue (win probability), red (lose amount, lose probability = 100 – probability win), and green (delay to trial outcome feedback) stimuli. Unlike the 2 × 2 experiment, participants could lose money and began with an account (5$) that could be deducted from. Spatial configurations of alternatives were be randomly presented in either horizontal rows or vertical columns. Corresponding attribute types were randomized on a trial-by-trial basis but arranged adjacently, allowing for equal ease of sampling with within-alternative or within-attribute fixation strategies. 150 trials were presented over the course of 3 separate blocks. In total, 30 dominated trials and 120 non-dominated trials were presented. Corresponding attribute types were randomized trial-by-trial but arranged adjacently, resulting in 48 possible four-alternative, four-attribute configurations (**Figure 4d**). Non-dominated trials in the 4 × 4 experiment were constructed using a Latin square design. Each alternative was assigned four of five possible attribute rankings, corresponding to the five possible magnitudes for each attribute (win amount: $.10, $.30, $.50, $.70, $.90; win probability: 10%, 30%, 50%, 70%, 90%; lose amount: −$.10, −$.30, −$.50, −$.70, −$.90; delay to feedback: 1s, 3s, 5s, 7s, 9s). Each non-dominated alternative is superior in one to three attributes and inferior in the remaining attributes to its paired alternative (see **Figure 2c** for an example choice menu). These alternatives were termed the amount win+, probability win+, amount lose+, and delay+ alternatives and were not mutually exclusive. Dominated trials in were constructed by ensuring that one attribute of the dominate alternative was superior and the remaining attributes were superior or equal in magnitude to the attributes of the other presented alternative.

### 4.4 Eye Tracking

#### RESULTS Choice behavior

##### Dominated trials

First, we evaluated the psychometric properties of participants’ choice behavior across all experiments. An analysis of two-alternative, two-attribute dominated trials showed that on average participants chose the optimal alternative 97% (SD = 3%) of the time. On four-alternative, four-attribute trials dominated trials, participants chose the optimal alternative 98% (SD = 3%) of the time on average. This finding provides evidence that participants can adequately identify the optimal alternative regardless of increased menu complexity.

##### Non-dominated trials

Analysis of two-alternative, two-attribute non-dominated trials revealed that participants chose the probability+ alternative (M = 67%, SD = 16%) significantly more often than the win+ alternative (M = 33%, SD = 16%), t(27) = 5.5573, p < 0.001.

Analysis of four-alternative, four-attribute non-dominated trials revealed that participants chose the probability+ alternative (M = 64%, SD = 19%) significantly more often than the win+ alternative (M = 23%, SD = 17%), t(21) = 5.5296, p < 0.001, the loss+ alternative (M = 9%, SD = 11%), t(21) = 9.6261, p < 0.001, and the delay+ alternative (M = 4%, SD = 4%), t(21) = 13.4392, p ¡ 0.001. Further, participants chose the win+ alternative significantly more often than the loss+ alternative, t(21) = 2.9919, p < 0.01, and the delay+ alternative t(21) = 5.5693, p < 0.001. Lastly, participants chose the loss+ alternative significantly more often than the delay+ alternative, t(21) = 2.3085, p < 0.05.

##### Expected value

In the 2×2 experiment win amount and win probability were the only two attributes. Expected value (EV) was calculated as: win amount * win probability. the 4×4 experiment consisted of two additional attributes: lose amount and delay to trial outcome feedback. In these experiments, expected value was calculated as: (win amount * probability win + lose amount * (1 – win probability)). Delay to trial outcome feedback had minimal influence on most participants’ choice behavior, so it was omitted from our expected value analyses.

Analysis of two-alternative, two-attribute non-dominated trials revealed that participants chose the greatest EV alternative (M = 66%, SD = 8%) significantly more often than the lowest EV alternative (M = 34%, SD = 8%), t(27) = 11.0503, p < 0.001. On two-alternative, four-attribute non-dominated trials, participants chose the greatest EV alternative (M = 73%, SD = 12%) significantly more often than the lowest EV alternative (M = 27%, SD = 12%), t(14) = 7.5361, p < 0.001. Four-alternative, two-attribute non-dominated trials each contained two equal EV pairs. Analysis of these trials revealed that participants did not choose the greatest EV alternatives (M = 57%, SD = 27%) significantly more often than the lowest EV alternatives (M = 43%, SD = 27%), t(14) = 0.9485, p = 0.36. On four-alternative, four-attribute non-dominated trials, participants chose the greatest EV alternative (M = 66%, SD = 11%) significantly more often than the second greatest EV alternative (M = 26%, SD = 8%), t(21) = 10.661, p < 0.001.

## Acknowledgements

We thank Hsin-Yi Hung for contributions to the Bayesian modeling effort. Support through NSF grant 1835202, NIH grant R01DA040990 (CRCNS), and from the U.S.-Israel Binational Science Foundation (CNCRS 2014612) is gratefully acknowledged.

## Appendix

### A Model equations and variables

For the convenience of the reader, we collect in this Appendix all variables used in the different models and for the computation of AIC, together with some key equations.

#### A.1 Expected Value

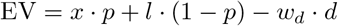

*x, p, l, d*: winning amount, winning probability, loss, delay

*w_d_*: weight for delay

#### A.2 Subjective Value

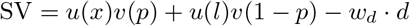

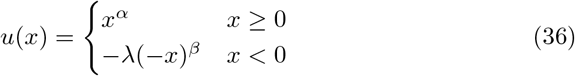

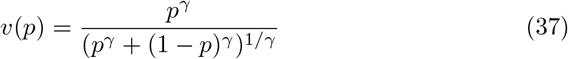

*α, β, λ, γ*: parameters for prospect theory

*u, v*: functions for prospect theory

#### A.3 Linear Combination

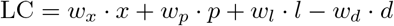

#### A.4 BI based on EV

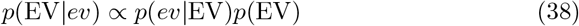

*σ*_EV_: standard deviation for the Gaussian distribution in likelihood function

#### A.5 BI based on SV

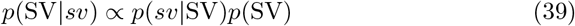

*σ*_SV_: standard deviation for the Gaussian distribution in likelihood function:

#### A.6 Latent Variable and prospect theory

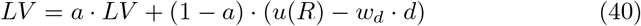

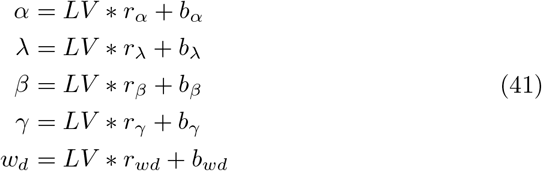

*a*: rate for the averaging of LV

*r*: rate of change

*b*: initial value for prospect theory parameters

#### A.7 aDDM

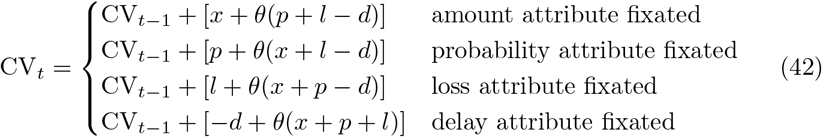

*t*: time

*θ*: parameter for those attributes not being fixated

#### A.8 DbS

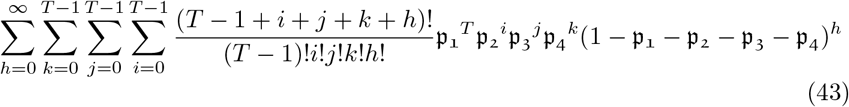

T: absolute threshold for accumulator

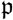: probability of having an increment

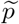: probability of sampling a comparison attribute within the immediate context

#### A.9 DbS with known attentional influence

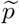: probability of sampling a comparison attribute within the immediate context

#### A.10 Two-layer leaky accumulator

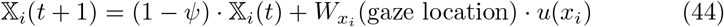

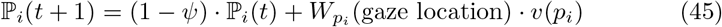

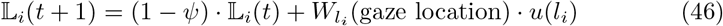

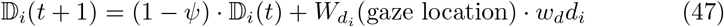

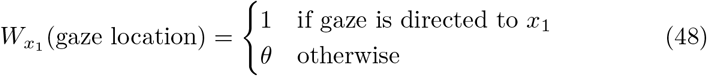

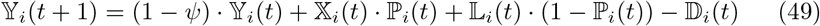

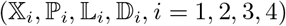: accumulators representing the activations of different attributes

*ψ*: degradation of information over time

*θ*: decreased influence of attribute when not given attention

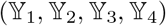: accumulators integrating the subjective values of the four options

#### A.11 PROMETHEE

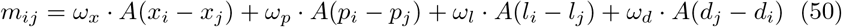

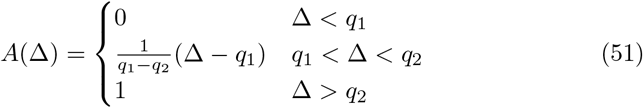

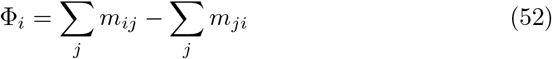

*m_ij_*: matrix element of multicriteria preference matrix

(*ω_x_, ω_p_, ω_l_, ω_d_*): weights for criteria

*q*1, *q*2: parameters determining the shape of the function *A*

Φ: net outranking flow

#### A.12 TN-pro

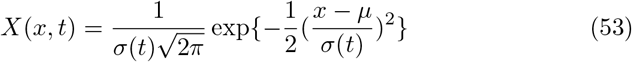

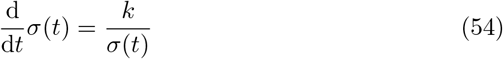

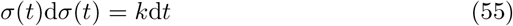

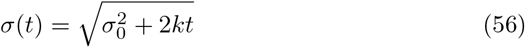

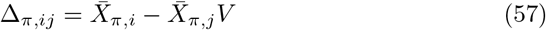

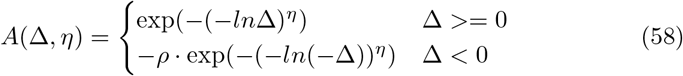

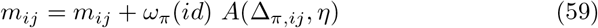

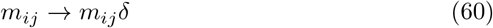

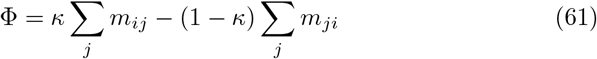

*X*: attribute value

*σ, μ*: std and center of the truncated Gaussian distribution

*k*: speed of forgetting

*σ*_0_: initial value for *σ*

*π*: attribute type

Δ: difference computed from pairwise comparison

*A, η, ρ*: advantage function and parameters controling its shape

*M, m*: advantage matrix and its elements

*ω, id*: the weight for different attribute type, the trial index

*δ*: leaky parameter for *M*

Φ, *κ*: outranking flow and its parameter

### B AIC and Likelihood

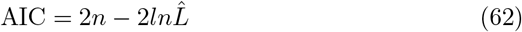

prob. for picking the option: 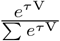

*n*: number of free parameters

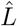: the model’s likelihood of data

V: final value for the option, could be EV, SV, LC

*τ*: fitting parameter used for getting prob.

### C Fitted parameters

This appendix shows the values of all parameters for all models, after fitting them using the maximum likelihood methods described in section 2.11.

#### C.1 Expected Value

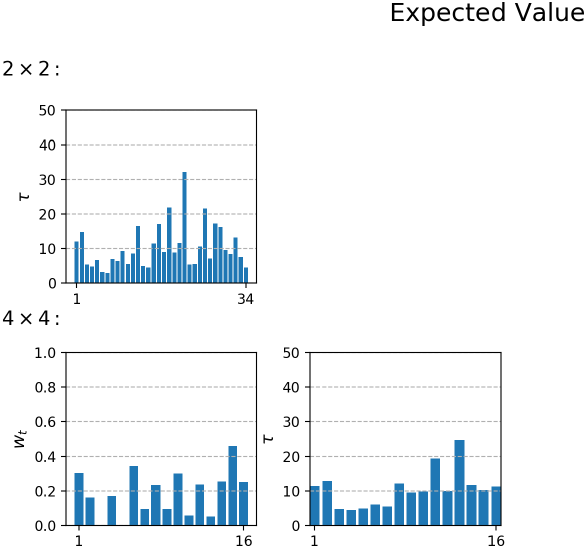

#### C.2 Subjective Value

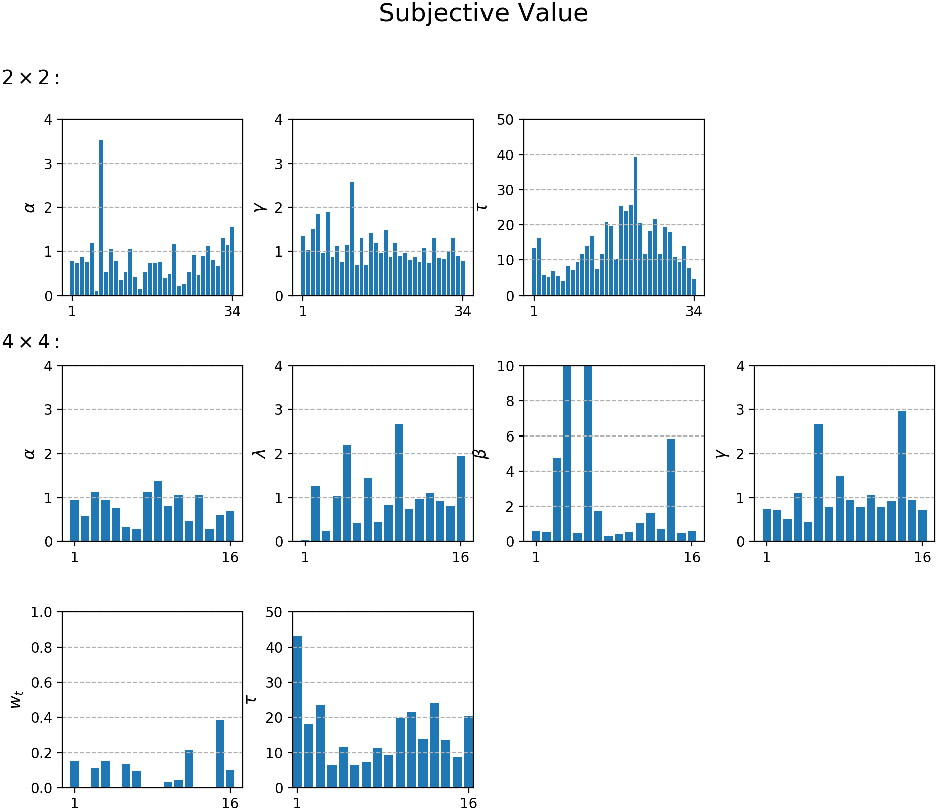

#### C.3 Linear Combination

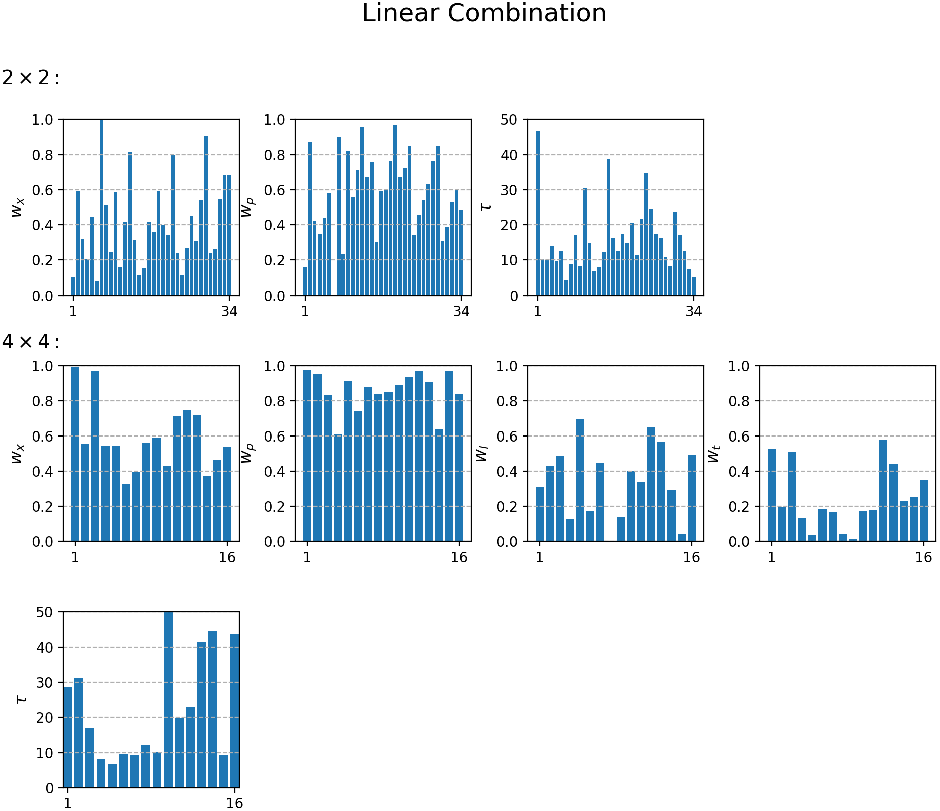

#### C.4 BI based on EV

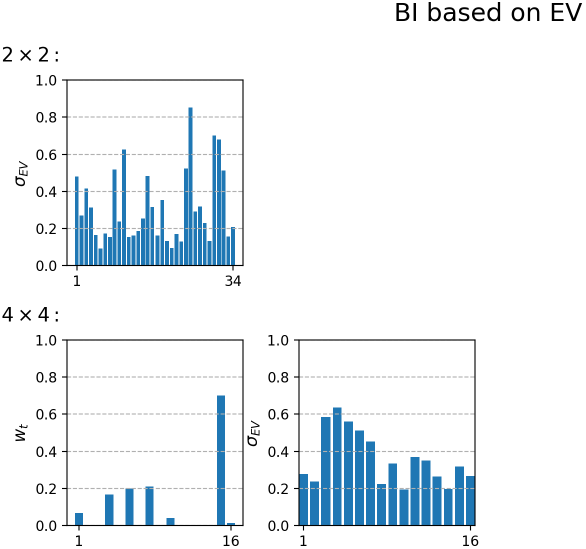

#### C.5 BI based on SV

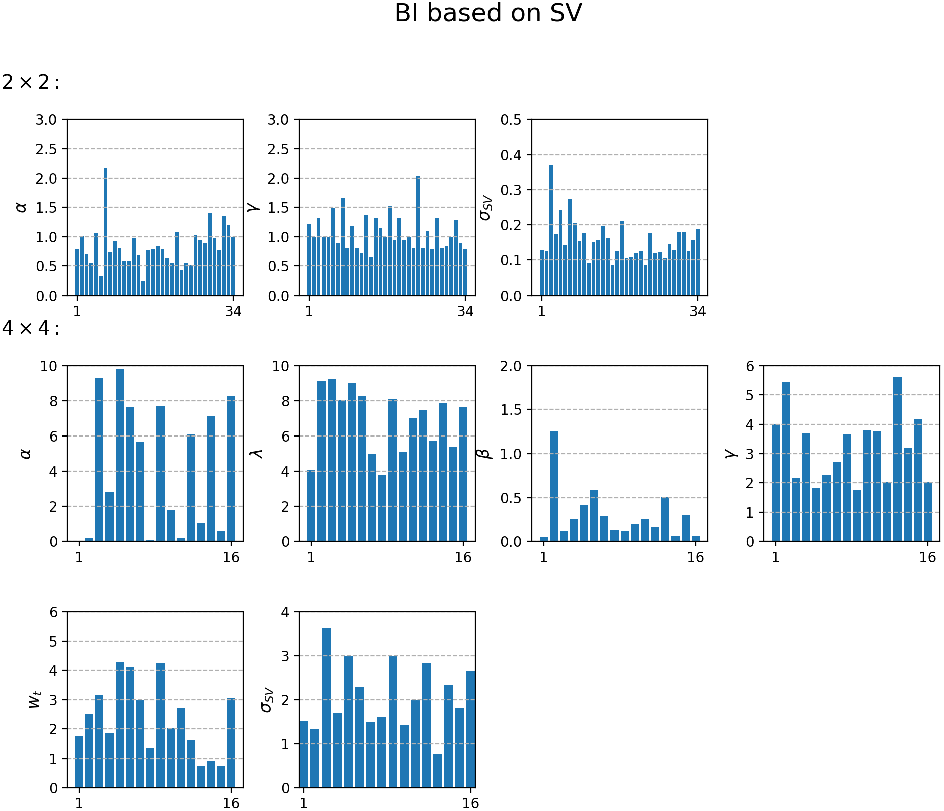

#### C.6 Latent Variable and prospect theory

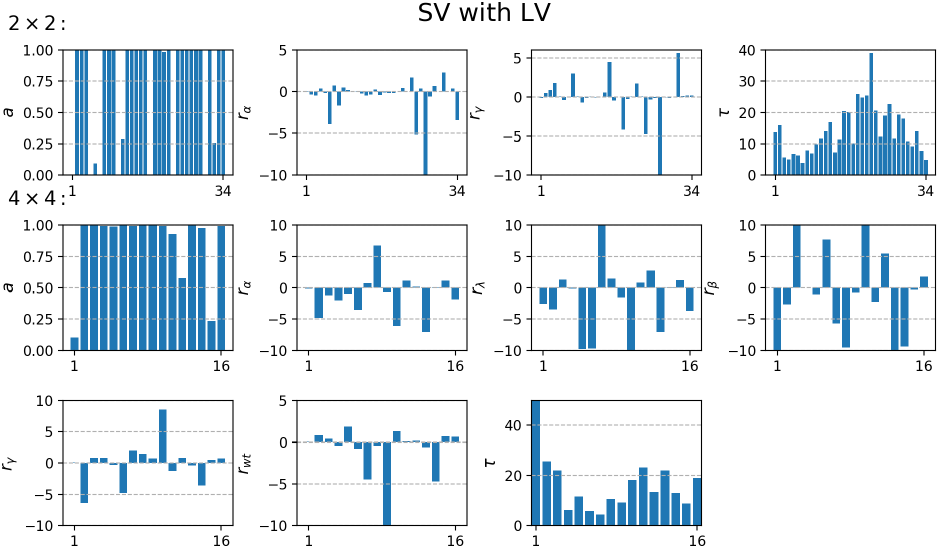

#### C.7 aDDM

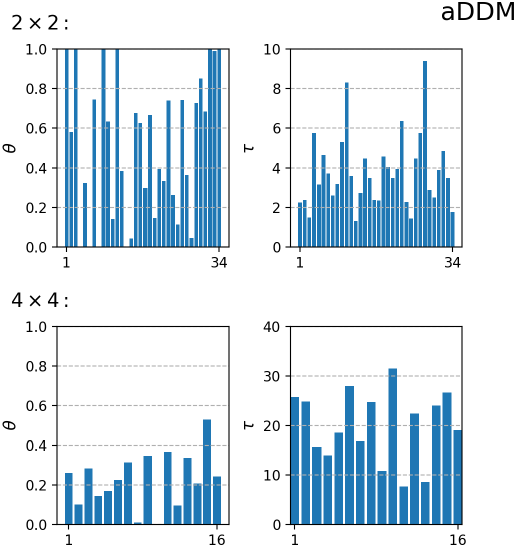

#### C.8 DbS

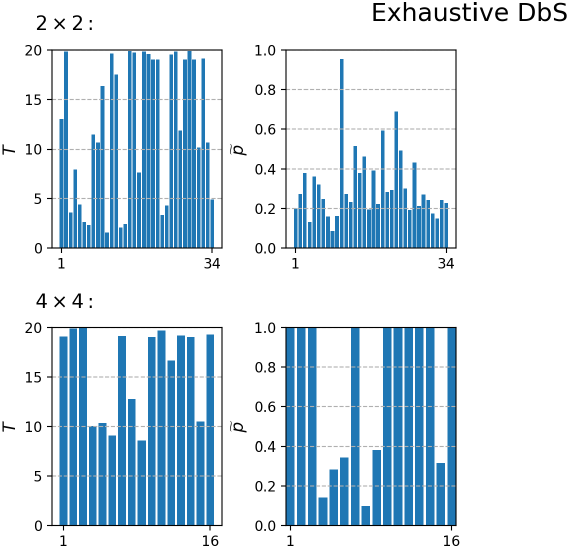

#### C.9 DbS with known attentional influence

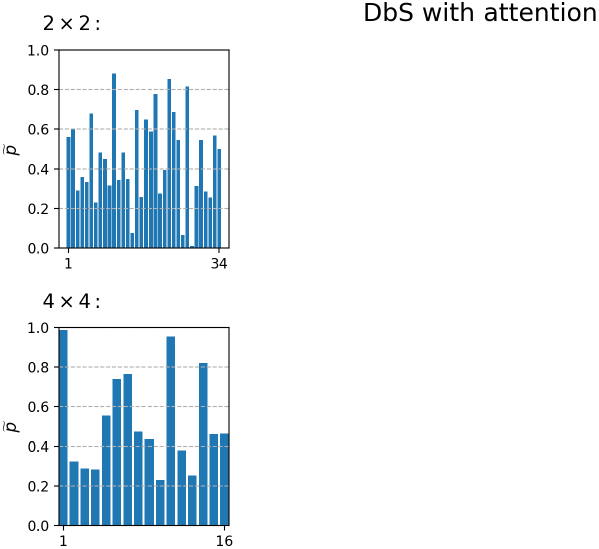

#### C.10 Two-layer leaky accumulator

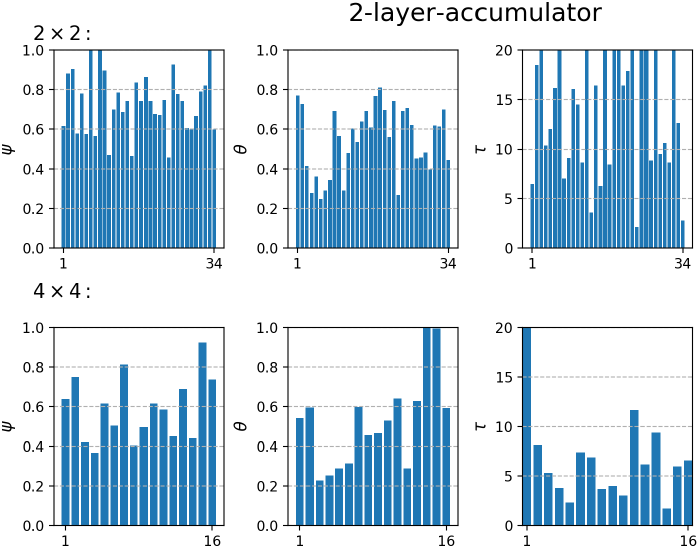

#### C.11 PROMETHEE

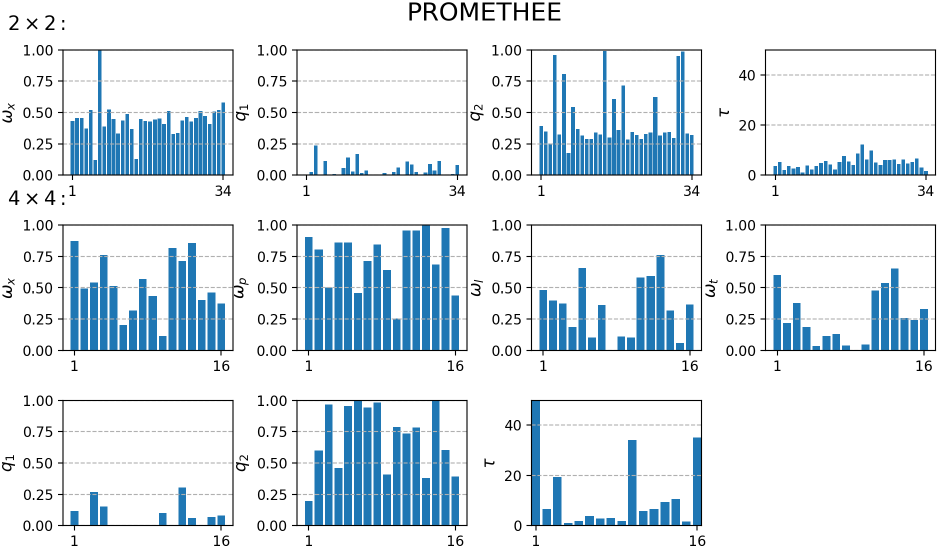

#### C.12 TNPRO

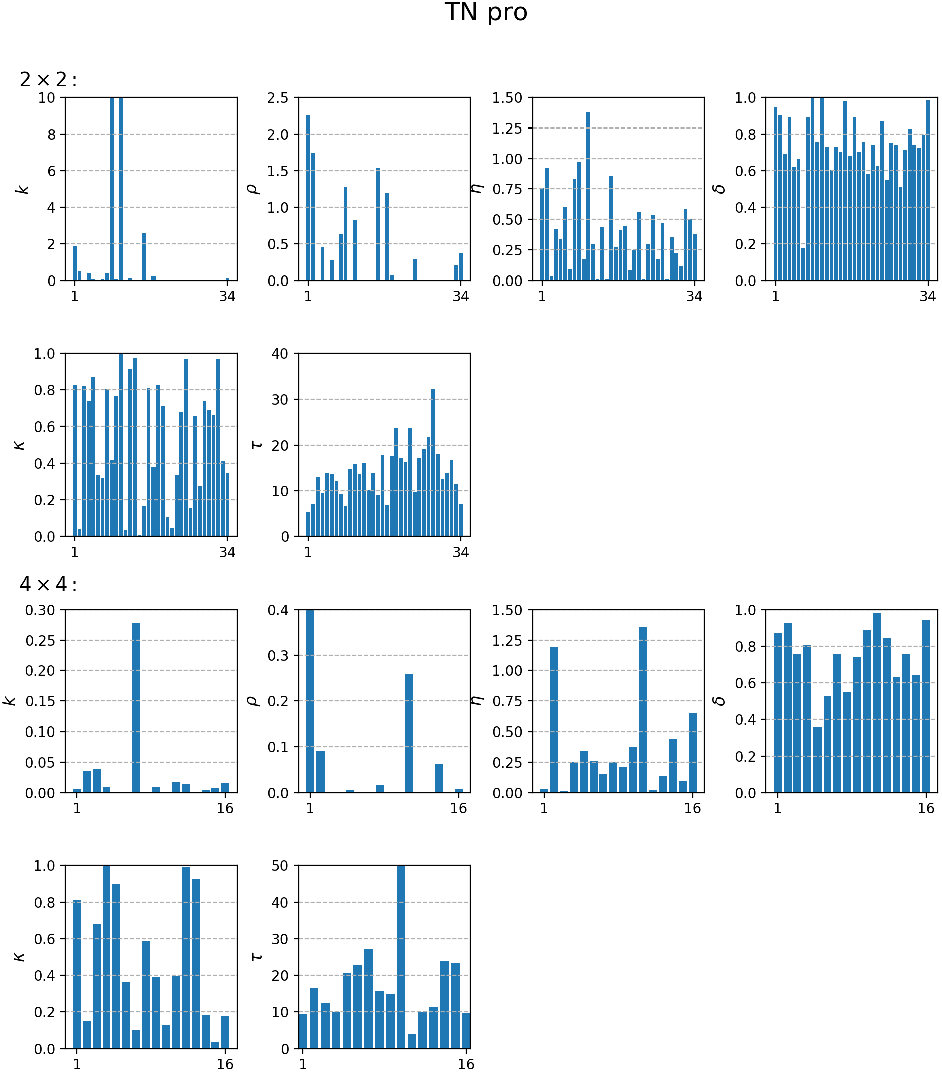

## Footnotes

1 This observation can be generalized to other indicators of attentional deployment: We have shown that predictions of computational models of covert attention [22, 14] correlate strongly not only with eye movements [26], as already mentioned, but also with the conscious selection of interesting parts of visual scenes identified by mouse clicks [21] or taps on the screen of an electronic tablet [15].

